# Uev1A counteracts oncogenic *Ras* stimuli in both polyploid and diploid cells

**DOI:** 10.1101/2025.04.17.649440

**Authors:** Qi Zhang, Yunfeng Wang, Xueli Fu, Ziguang Wang, Yang Zhang, Lizhong Yan, Yuejia Wang, Muhan Yang, Dongze Song, Ruixing Zhang, Hongru Zhang, Shian Wu, Shaowei Zhao

**Author notes:** These authors contributed equally (Qi Zhang, Yunfeng Wang, Xueli Fu, Ziguang Wang).

## Abstract

Oncogenic *Ras* is known to induce DNA replication stress, leading to cellular senescence or death. In contrast, we found that it can also trigger polyploid *Drosophila* ovarian nurse cells to die by inducing aberrant division stress. To explore intrinsic protective mechanisms against this specific form of cellular stress, here we conducted a genome-wide genetic screen and identified the E2 enzyme Uev1A as a key protector. Reducing its expression levels exacerbates the nurse cell death induced by oncogenic *Ras*, while overexpressing it or its human homologs, UBE2V1 and UBE2V2, mitigates this effect. Although Uev1A is primarily known for its non-proteolytic functions, our studies demonstrate that it collaborates with the E3 APC/C complex to mediate the proteasomal degradation of Cyclin A, a key cyclin that drives cell division. Furthermore, Uev1A and UBE2V1/2 also counteract oncogenic *Ras*-driven tumorigenesis in diploid cells, suppressing the overgrowth of germline tumors in *Drosophila* and human colorectal tumor xenografts in nude mice, respectively. Remarkably, elevated expression levels of UBE2V1/2 correlate with improved survival rates in human colorectal cancer patients harboring oncogenic *KRAS* mutations, indicating that their upregulation could represent a promising therapeutic strategy.

## Introduction

Although the human body contains trillions of cells that could potentially be targeted by oncogenic mutations, cancers arise infrequently throughout a human lifetime, suggesting the presence of effective resistance mechanisms (***Lowe et al., 2004***). One key mechanism involves the induction of DNA replication stress by these oncogenic mutations, leading to cellular senescence or death (***Hills and Diffley, 2014; Kotsantis et al., 2018***). This mechanism is mediated through the activation of the DNA damage response (DDR) pathway and the tumor suppressor protein p53, which are triggered by DNA double-strand breaks (DSBs). Only the cells that escape senescence and death, such as those harboring *p53* mutations, can undergo transformation by oncogenic mutations, thereby initiating tumorigenesis **(*Gaillard et al., 2015; Igarashi et al., 2024; Macheret and Halazonetis, 2015***). Among the most frequently mutated oncogenes in human cancers are the *RAS* genes (*KRAS*, *NRAS*, and *HRAS*), which mutations are present in 20-30% of all cancer cases (***Gimple and Wang, 2019; Sanchez-Vega et al., 2018***). These mutations often occur at codons 12, 13, and 61, resulting in RAS small GTPases being locked in a constitutively active, GTP-bound state **(*Moore et al., 2020***). Notably, previous studies have shown that oncogenic *HRAS^G12V^* can induce DNA replication stress via the DDR pathway, ultimately leading to cellular senescence **(*Di Micco et al., 2006; Serrano et al., 1997*)**.

The *Ras oncogene at 85D* gene (*Ras85D*, hereafter referred to as *Ras*) in *Drosophila* exhibits high homology to human *RAS* genes (***Neuman-Silberberg et al., 1984***). Our previous research demonstrated that oncogenic *Ras^G12V^* triggers cell death in *Drosophila* ovarian nurse cells (***Zhang et al., 2024***), a type of post-mitotic germ cells that undergo G/S endoreplication to become polyploid (***Hammond and Laird, 1985***) (***Figure 1A***). Notably, oncogenic *Ras^G12V^* promotes the division of mitotic germ cells, the precursors to nurse cells. Furthermore, monoallelic deletion of the *cyclin A* (*cycA*) or *cyclin-dependent kinase 1* (*cdk1*) gene (*cycA^+/-^* or *cdk1^+/-^*) suppresses *Ras^G12V^*-induced nurse cell death (***Zhang et al., 2024***). While CycA, a key cyclin promoting cell division, is not expressed in normal nurse cells **(*Lilly et al., 2000; Lilly and Spradling, 1996***), its ectopic expression in nurse cells can trigger their death (as observed in this study). These findings suggest that the nurse cell death induced by oncogenic *Ras^G12V^* is primarily due to aberrant promotion of cell division. *Drosophila* ovarian nurse cells thus offer a valuable model for studying this specific form of cellular stress.

**Figure 1.**
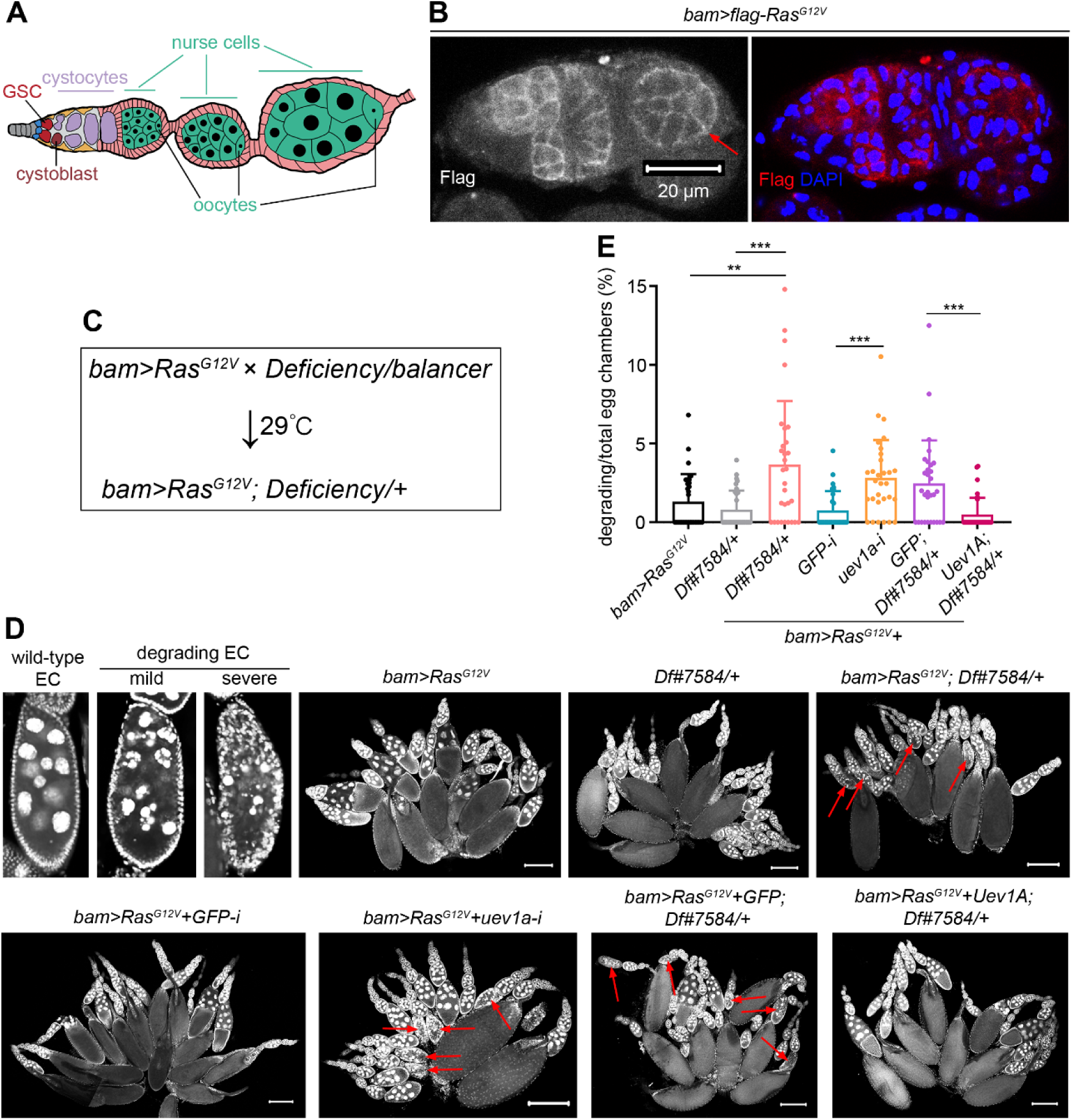
Genetic screen identifies Uev1A as a crucial protector against *Ras^G12V^*-induced nurse cell death. (A) Schematic cartoon for *Drosophila* ovariole. During oogenesis, GSC undergoes asymmetric division to generate two daughter cells: one that self-renews and maintains GSC identity, and the other, called a cystoblast, that differentiates to support oogenesis. As differentiation progresses, each cystoblast performs four rounds of division with incomplete cytokinesis to produce 16 interconnected cystocytes, establishing a germline cyst. This germline cyst is then surrounded by epithelial follicle cells to form an egg chamber. Within each egg chamber, one of the 16 germ cells becomes the oocyte, while the remaining 15 differentiate into nurse cells. These nurse cells undergo G/S endocycling, becoming polyploid to aid in oocyte development. (B) Representative germarium and early-stage egg chamber with Flag-Ras^G12V^ overexpression driven by *bam-GAL4-VP16*. The red arrow denotes an early-stage egg chamber. (C) Genetic screening strategy. Genotype of “*bam>Ras^G12V^*”: *bam-GAL4-VP16/FM7;; UASp-Ras^G12V^/TM6B*. (D) Representative ovaries and egg chambers (DAPI staining). The red arrows in (D) denote degrading egg chambers. Scale bars: 200 μm. (E) Quantification data. 30 ovaries from 7-day-old flies were quantified for each genotype. Statistical significance was determined using t test (groups = 2) or one-way ANOVA (groups > 2): ** (*P* < 0.01) and *** (*P* < 0.001). The online version of this article includes the following source data for figure 1: **Source data 1.** Screen results. **Source data 2.** All genotypes. **Source data 3.** Raw quantification data.

Ubiquitination not only mediates protein degradation through the proteasome or lysosome but also plays crucial regulatory roles in various biological processes **(*Dikic, 2017; Kwon and Ciechanover, 2017***). These functions are largely determined by the topological structures of the polyubiquitin chains, where ubiquitin can be linked to another ubiquitin via any of its seven lysine (K) residues or its first methionine residue. Among these, K11- and K48-linked polyubiquitin primarily mediates the proteasomal degradation. In contrast, K63 linkage facilitates the lysosomal degradation of protein aggregates and damaged organelles through autophagy. Additionally, K63 linkage is involved in non-degradative processes such as DNA repair, kinase activation, and protein transport **(*Kwon and Ciechanover, 2017***). Ubc13 is a ubiquitin-conjugating (E2) enzyme that specifically catalyzes the formation of the K63 linkage. However, this activity requires a unique cofactor known as Ubc variant (Uev). Uev proteins lack the catalytic cysteine residue necessary for ubiquitin thioester formation; instead, they work with Ubc13 to form a functional E2 complex (***Hofmann and Pickart, 1999; McKenna et al., 2001***). These E2 enzymes have been shown to play regulatory roles in various biological processes, including cell death (***Ma et al., 2013***), DNA repair (***Broomfield et al., 1998***), DDR, and innate immunity (***Andersen et al., 2005; Bai et al., 2020; Zhou et al., 2005***). However, it still remains unknown whether they also possess proteasomal proteolytic functions and whether such functions hold significant biological roles.

In this study, to investigate the intrinsic protective mechanisms against *Ras^G12V^*-induced nurse cell death in *Drosophila*, we conducted a genome-wide genetic screen and identified Uev1A as a crucial protector. Mechanistically, Uev1A works in conjunction with the anaphase-promoting complex or cyclosome (APC/C) to degrade the essential cyclin CycA via the proteasome, highlighting the critical role of its proteolytic function. Additionally, Uev1A and its human homologs, UBE2V1 and UBE2V2, also counteract oncogenic *Ras*-driven tumorigenesis in diploid cells, inhibiting the overgrowth of *Drosophila* germline tumors and human colorectal tumor xenografts in nude mice, respectively. These findings suggest that upregulation of UBE2V1/2 could represent a promising therapeutic approach for human colorectal cancers with *RAS* mutations.

## Results

### Genetic screen identifies Uev1A as a crucial protector against *Ras^G12V^*-induced nurse cell death

While oncogenic *Ras^G12V^* triggers cell death in nurse cells, it does not have the same effect in cystocytes, the precursor cells before differentiating into nurse cells (***Zhang et al., 2024***). To confirm Ras^G12V^ expression in cystocytes, we generated a *UASz-flag-Ras^G12V^* transgenic fly strain, which phenocopied the effects observed with the previously used *UASp-Ras^G12V^*. RAS small GTPases need to be anchored to the inner cell membrane for their proper function (***Willingham et al., 1980***). Using *bam-GAL4-VP16* (***Chen and McKearin, 2003***), we overexpressed Ras^G12V^ specifically in cystocytes (*bam>flag-Ras^G12V^*) at 29°C and observed membrane-localized Flag signals, validating the normal expression of Ras^G12V^ (***Figure 1B***). Also, Flag signals were detected in some early-stage nurse cells, although these signals gradually diminished over time (***Figure 1B***). During cell death, nurse cell nuclei progress through distinct morphological stages: from large and round, to disorganized and condensed, and finally to completely fragmented into small, spherical structures (***Figure 1D***). Given the interconnection and synchronous death of all nurse cells within an egg chamber, we quantified this death phenotype at the egg-chamber level. Notably, nurse cell death remained very low in *bam>flag-Ras^G12V^* fly ovaries (***Figure 1D, E***). This may be attributed to either insufficient levels of the oncoproteins or the presence of a protective mechanism.

To explore the potential protective mechanism, we conducted a genetic modifier screen by introducing individual genome-wide *Deficiencies* (361 lines) into *bam>Ras^G12V^*flies (***bam>Ras^G12V^; Deficiency/+, Figure 1C***). Of note, ovaries from *bam>Ras^G12V^; Df#7584/+* flies exhibited the highest incidence of degrading egg chambers with dying nurse cells per ovary (**see Source data 1**). One gene deleted in this deficiency line is *uev1a*, and RNAi targeting *uev1a* reproduced the phenotype seen with *Df#7584* (***Figure 1D, E***). In this and all subsequent experiments, we quantified the nurse-cell-death phenotype using the percentage of degrading to total egg chambers per ovary (***Figure 1E***), a method that is more precise than our approach in the genetic screen. To further verify the role of *uev1a*, we generated a *UASz-uev1a* transgenic fly strain. Overexpression of Uev1A successfully rescued the nurse-cell-death phenotype in *bam>Ras^G12V^; Df#7584/+* fly ovaries (***Figure 1D, E***), confirming that Uev1A is a crucial protector against *Ras^G12V^*-induced nurse cell death.

### Protective role of Uev1A against the nurse cell death induced by direct overexpression of Ras^G12V^

In contrast to the scenario above (*bam>Ras^G12V^*), direct overexpression of Ras^G12V^ in nurse cells using *nos-GAL4-VP16* (Rørth, 1998; Van Doren et al., 1998) (*nos>Ras^G12V^*) resulted in substantial cell death even at 25°C (***Figure 2A, D***) (***Zhang et al., 2024***). Such observation prompted us to investigate the role of Uev1A in this context. Notably, the incidence of dying nurse cells was markedly elevated in *nos>Ras^G12V^+uev1a-RNAi* ovaries compared to the *nos>Ras^G12V^+GFP-RNAi* control ovaries (***Figure 2A, D***). To further validate this, we generated two *uev1a* mutants using the CRISPR/Cas9 technique: *uev1a*^Δ*1*^ and *uev1a*^Δ*2*^. Both mutations consist of small deletions that cause frame shifts within the second coding exon of the *uev1a* gene (***Figure 2B***). Of note, flies with the *uev1a*^Δ*1/*Δ*1*^, *uev1a*^Δ*1/*Δ*2*^, *uev1a*^Δ*1*^*/Df#7584*, *uev1a*^Δ*2/*Δ*2*^, or *uev1a*^Δ*2*^*/Df#7584* genotype could not survive into adults, suggesting that both mutations are strong loss-of-function alleles. Consistent with the effects observed with *uev1a-RNAi*, either mutation markedly increased the incidence of dying nurse cells in *nos>Ras^G12V^* ovaries (***Figure 2C, D***).

**Figure 2.**
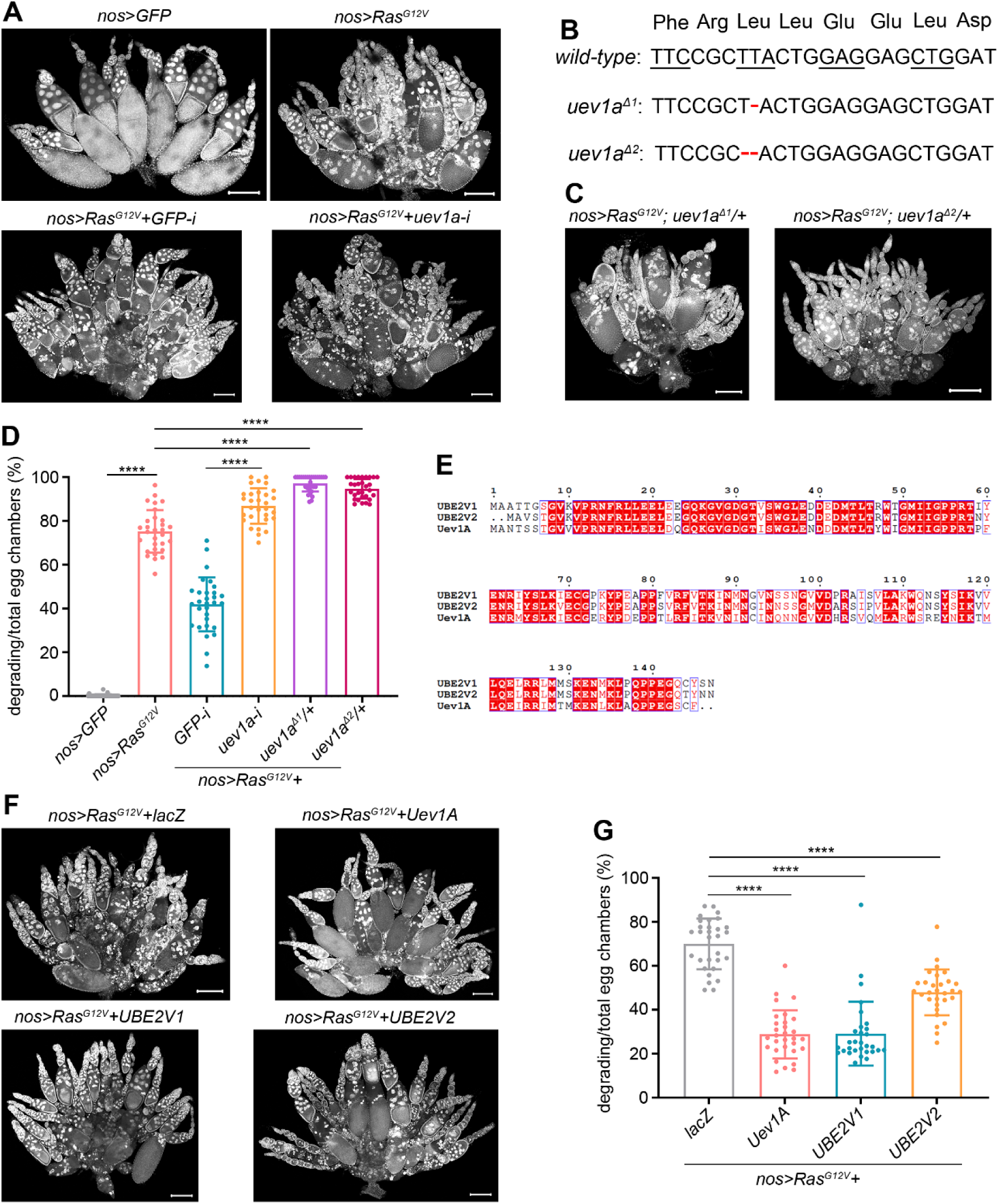
Uev1A protects against the nurse cell death induced by direct overexpression of Ras^G12V^. (A, C, and F) Representative samples (DAPI staining). (B) Molecular information of the *uev1a*^Δ*1*^ and *uev1a*^Δ*2*^ mutations. The red dashed lines represent nucleotide deletions. (D and G) Quantification data. 30 ovaries from 3-day-old flies were quantified for each genotype. Statistical significance was determined using t test (groups = 2) or one-way ANOVA (groups > 2): **** (*P* < 0.0001). (E) Protein sequence alignment of Uev1A, UBE2V1, and UBE2V2. It was performed using CLUSTALW and ESPript 3.0 software. The online version of this article includes the following source data and figure supplement for figure 2: **Source data 2.** All genotypes. **Source data 3.** Raw quantification data. **Figure supplement 1.** Uev1A protects against *Yki^3SA^*-induced nurse cell death.

We next investigated whether upregulating Uev1A could mitigate the nurse cell death induced by direct overexpression of *Ras^G12V^*. Remarkably, compared to the *nos>Ras^G12V^+lacZ* control ovaries, the incidence of dying nurse cells was significantly reduced in *nos>Ras^G12V^+Uev1A* ovaries (***Figure 2F, G***). Uev1A has two human homologs, UBE2V1 and UBE2V2, which share 67% (85%) and 67% (86%) sequence identities (similarities) with Uev1A, respectively (***Figure 2E***). We generated transgenic fly strains for both *UASz-UBE2V1* and *UASz-UBE2V2*. Notably, overexpression of either UBE2V1 or UBE2V2 significantly mitigated *Ras^G12V^*-induced nurse cell death, similar to the effects observed with Uev1A overexpression (***Figure 2F, G***). Taken together, these results further confirm the protective role of Uev1A against *Ras^G12V^*-induced nurse cell death.

Then, we were intrigued by the potential of Uev1A to protect against the nurse cell death induced by other oncogenic mutations. Yorkie (Yki), a key oncoprotein in the Hippo pathway (***Huang et al., 2005***), has a hyperactive form known as Yki^3SA^ **(*Oh and Irvine, 2009***). Its overexpression using *nos-GAL4-VP16* (*nos>Yki^3SA^*) at 29°C, but not at 25°C, could induce nurse cell death (***Figure 2—figure supplement 1***), albeit much less severe than that induced by *nos>Ras^G12V^* at 25°C (compared with ***Figure 2A, D***). Of note, the nurse cell death induced by oncogenic *Yki^3SA^* was significantly alleviated by Uev1A overexpression (***Figure 2—figure supplement 1***), implying a broad role of Uev1A in this process. However, due to the mild effect of Yki^3SA^ on triggering nurse cell death, we focused on Ras^G12V^ in our subsequent studies.

### The DDR pathway and p53 play opposite roles in *Ras^G12V^*-induced nurse cell death

To investigate how Uev1A protects against *Ras^G12V^*-induced nurse cell death, we first sought to further explore the mechanisms driving this cell death. Since egg chambers contain both nurse cells and oocytes (***Figure 1A***), we considered the possibility that the death signal could originate from oocytes. The *Bicaudal D* (*BicD*) gene is essential for oocyte determination, and egg chambers deficient in it fail to specify oocytes (***Suter and Steward, 1991***). We confirmed this by using *nos>BicD-RNAi* (***Figure 3—figure supplement 1A***). Importantly, nurse cell death was much more pronounced in *nos>BicD-RNAi+Ras^G12V^* egg chambers than in *nos>BicD-RNAi+GFP* control ones (***Figure 3—figure supplement 1A, B***), indicating that the death signal is intrinsic to nurse cells rather than originating from oocytes.

**Figure 3.**
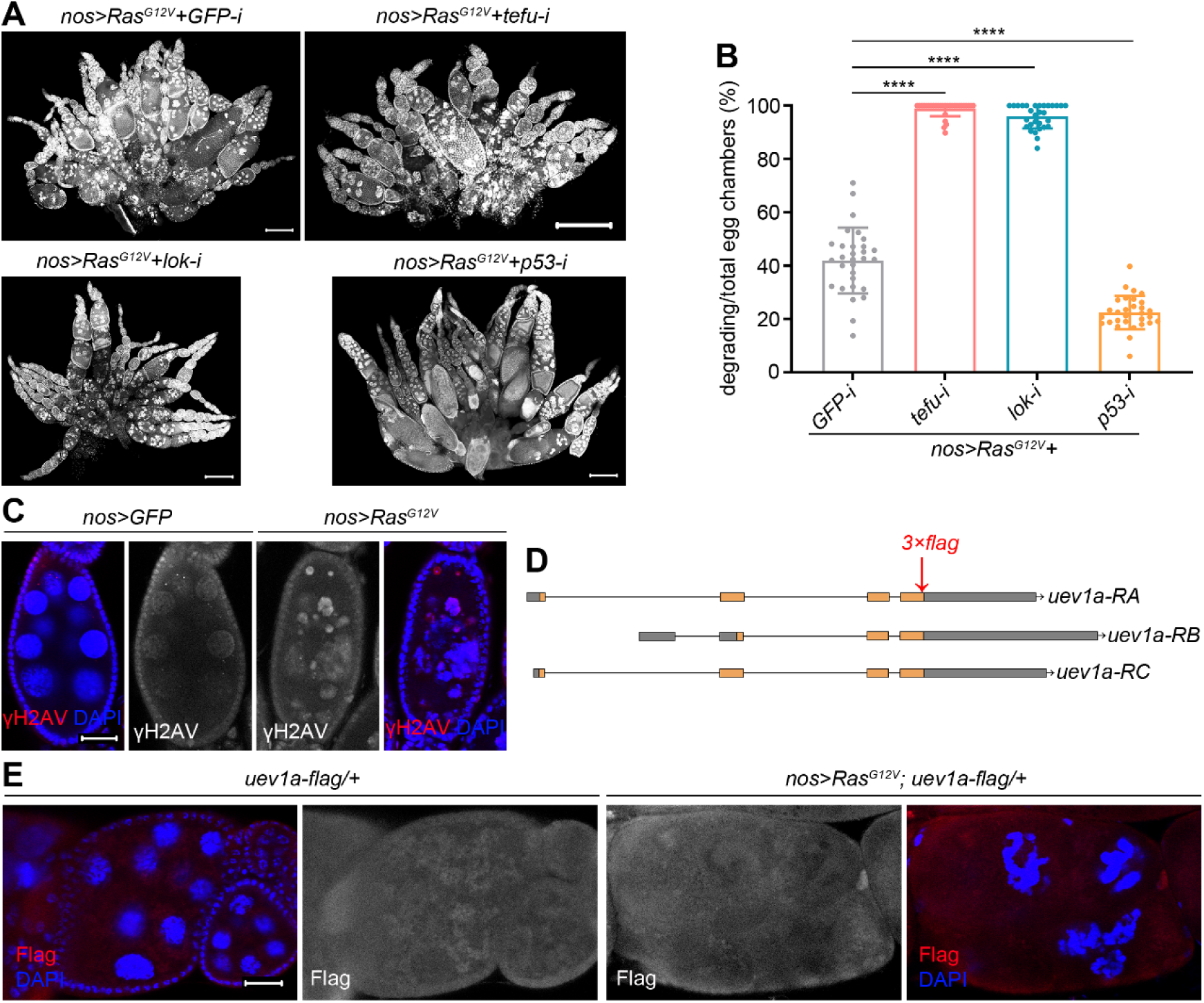
Roles of the DDR pathway and p53 in *Ras^G12V^*-induced nurse cell death. (A) Representative ovaries (DAPI staining). Scale bars: 200 μm. (B) Quantification data. 30 ovaries from 3-day-old flies were quantified for each genotype. Statistical significance was determined using one-way ANOVA: **** (*P* < 0.0001). (C and E) Representative samples. Scale bars: 20 μm. (D) Schematic cartoon for *uev1a-flag* knock-in. The online version of this article includes the following source data and figure supplement for figure 3: **Source data 2.** All genotypes. **Source data 3.** Raw quantification data. **Figure supplement 1.** Oncogenic *Ras^G12V^* intrinsically triggers nurse cell death. **Figure supplement 2.** Uev1A is expressed in stretch follicle cells. **Figure supplement 3.** Uev1A does not directly degrade the Ras^G12V^ oncoproteins.

Oncogenic *RAS* can induce DNA replication stress in diploid cells, thereby activating the DDR pathway and p53 to trigger cellular senescence or death **(*Di Micco et al., 2006; Serrano et al., 1997***). To investigate whether DDR plays a similar role in *Ras^G12V^*-induced nurse cell death, we targeted two key DDR genes: *telomere fusion* (*tefu*, encoding *Drosophila* ATM) **(*Oikemus et al., 2004; Silva et al., 2004; Song et al., 2004***) and *loki* (*lok*, encoding *Drosophila* Chk2) (***Masrouha et al., 2003; Xu et al., 2001***). Strikingly, knockdown of either gene markedly enhanced this cell death (***Figure 3A, B***), revealing a protective role for the DDR pathway. By contrast, knockdown of *p53* suppressed this cell death (***Figure 3A, B***), demonstrating opposing functions for DDR and p53 in this context. To assess DNA DSBs and the ensuing DDR, we monitored the phosphorylation of γH2AV, the *Drosophila* histone variant analogous to mammalian H2AX (***Lake et al., 2013***). As egg chamber developmental stages are difficult to discern during degradation, we compared size-matched egg chambers, which are typically stage-matched under normal conditions and have comparable antibody penetration. Elevated γH2AV staining was observed in degrading *nos>Ras^G12V^* egg chambers compared to *nos>GFP* controls (***Figure 3C***), indicating a heightened burden of DNA DSBs.

Given that RNAi targeting *tefu* or *lok* phenocopied that of *uev1a*, we hypothesized that Uev1A protects against *Ras^G12V^*-induced nurse cell death through the DDR pathway. If so, Uev1A may also be upregulated in response to *Ras^G12V^*-driven stress. To test this, we initially attempted to generate an antibody against Uev1A using its full protein sequence as the antigen; however, this approach was unsuccessful. As an alternative, we created a *uev1a-flag* knock-in fly strain by inserting the coding sequence of a 3xFlag tag immediately before the stop codon (“TAG”) of the *uev1a* gene (***Figure 3D***). These knock-in flies were homozygous viable and fertile, indicating that the Flag insertion did not disrupt Uev1A’s normal function. Previous studies have shown that Uev1A can activate JNK signaling (***Ma et al., 2013***), which acts in the surrounding follicle cells to promote nurse cell removal during late oogenesis **(*Timmons et al., 2016***). Indeed, we detected Uev1A-Flag signals in such follicle cells (***Figure 3—figure supplement 2***), confirming that these signals reflect endogenous Uev1a expression in *Drosophila* ovaries. Surprisingly, Uev1A-Flag signals were low in both *nos>Ras^G12V^; uev1a-flag/+* and *uev1a-flag/+* control nurse cells, with no significant differences between the two groups (***Figure 3E***). This suggests that Uev1A expression remains at basal levels, rather than being upregulated by *Ras^G12V^*-driven stress in this context. Although we cannot rule out the possibility that Uev1A may play a role in the DDR pathway at these basal levels, these findings prompted us to explore its potential proteolytic function as an E2 enzyme.

Following the Occam’s Razor principle, we first investigated whether Uev1A is involved in downregulating Ras^G12V^ oncoprotein levels. To address the challenges of analyzing membrane-localized signals in *nos>flag-Ras^G12V^*nurse cells due to severe cell death, we performed this assay in *bam>flag-Ras^G12V^*ovaries at 29°C. Notably, similar membrane-localized Flag signals were detected in both *bam>flag-Ras^G12V^+Uev1A* and *bam>flag-Ras^G12V^+GFP* control germ cells, including nurse cells (***Figure 3—figure supplement 3***). This result suggests that Uev1A does not downregulate Ras^G12V^ oncoprotein levels.

### Uev1A downregulates CycA protein levels in *Ras^G12V^*-induced dying nurse cells

Our previous research demonstrated that oncogenic *Ras^G12V^*promotes germline stem cell (GSC) over-proliferation by activating the mitogen-activated protein kinase (MAPK) pathway (***Zhang et al., 2024***). This finding led us to investigate whether the same pathway influences *Ras^G12V^*-induced nurse cell death. Notably, knockdown of *downstream of raf1* (*dsor1*, encoding *Drosophila* MAPKK) **(*Tsuda et al., 1993***) or *rolled* (*rl*, encoding *Drosophila* MAPK) (***Biggs and Zipursky, 1992***) significantly mitigated *Ras^G12V^*-induced nurse cell death (***Figure 4A, B***), highlighting the pivotal role of the MAPK pathway in this process.

**Figure 4.**
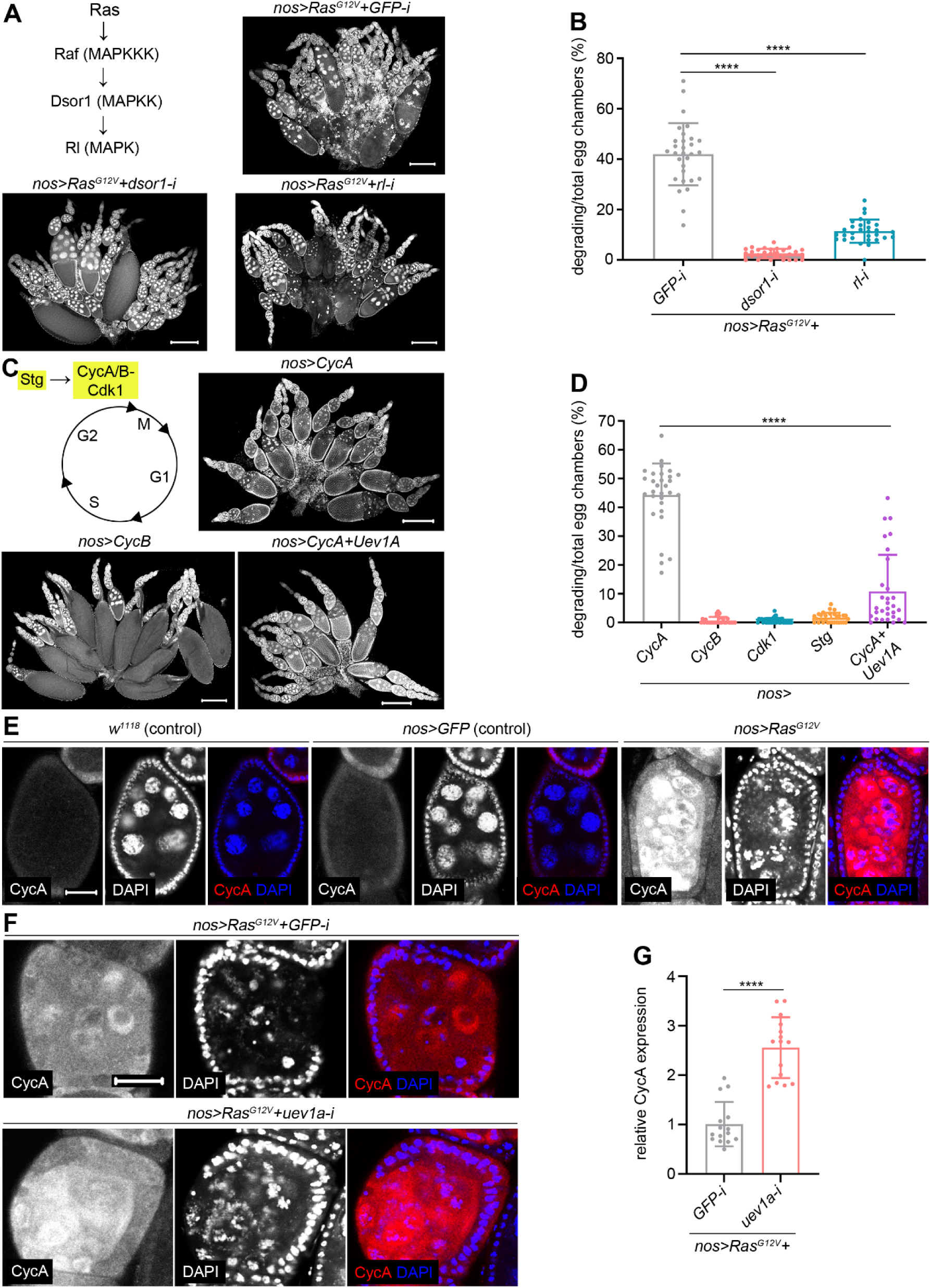
Uev1A collaborates with CycA to mitigate *Ras^G12V^*-induced nurse cell death. (A, C, E, and F) Representative ovaries. DAPI staining in (A and C). Scale bars: 200 μm in (A and C), 20 μm in (E and F). (B, D, and G) Quantification data. 30 ovaries (B and D) and 15 size-matched egg chambers (G) from 3-day-old flies were quantified for each genotype. Statistical significance was determined using t test (groups = 2) or one-way ANOVA (groups > 2): **** (*P* < 0.0001). The online version of this article includes the following source data for figure 4: **Source data 2.** All genotypes. **Source data 3.** Raw quantification data.

The Ras/MAPK pathway is well known to promote cell cycle progression **(*Gimple and Wang, 2019***). Our previous work demonstrated that reducing the gene dosage of *CycA* or *Cdk1* suppresses *Ras^G12V^*-induced nurse cell death (***Zhang et al., 2024***). In addition to CycA and Cdk1, Cyclin B (CycB) and String (Stg) are two other essential regulators that drive cell division (***Figure 4C***) (***Edgar and Lehner, 1996***). To determine if individual overexpression of these regulators could induce nurse cell death, we tested each and found that only CycA triggered this phenotype (***Figure 4C, D***). Notably, this cell death was suppressed by co-overexpression of CycA and Uev1A (***Figure 4C, D***), indicating a genetic interaction between them. In line with previous findings (***Lilly et al., 2000; Lilly and Spradling, 1996***), CycA protein was undetectable in wild-type endocycling nurse cells (***Figure 4E***). In stark contrast, it was abundantly expressed in dying *nos>Ras^G12V^*nurse cells (***Figure 4E***), underscoring a critical role for CycA in this cell death process. Additionally, we assessed CycA protein levels in size-matched *nos>Ras^G12V^* nurse cells under either *uev1a* or *GFP* (control) knockdown condition. Notably, *uev1a* knockdown increased CycA levels compared with the controls (***Figure 4F, G***), demonstrating that Uev1A downregulates CycA protein levels in *Ras^G12V^*-induced dying nurse cells.

### Uev1A collaborates with the APC/C complex to mitigate *Ras^G12V^*-induced nurse cell death

It is well-established that the APC/C complex primarily functions as an E3 ligase to facilitate the degradation of CycA during cell cycle progression (***Sudakin et al., 1995***). Thus, a compelling model to explain our findings is that Uev1A collaborates with the APC/C complex to degrade CycA. Cell division cycle 27 (Cdc27, *Drosophila* APC3) is an essential part of the substrate recognition TPR lobe within the APC/C complex (***Yamano, 2019***). In *bam>Ras^G12V^+cdc27-RNAi* ovaries, we observed dying nurse cells, a phenotype that was exacerbated with mutations in either *uev1a*^Δ*1*^ or *uev1a*^Δ*2*^ (***Figure 5A, B***). Furthermore, knocking down *cdc27* could increase the incidence of dying nurse cells in *bam>Ras^G12V^+uev1a-RNAi* ovaries (***Figure 5A, B***). Also, we investigated Fizzy-related (Fzr), the *Drosophila* homolog of Cdh1, that is a critical activator of APC/C-dependent proteolysis. It is known that Fzr functions to downregulate mitotic cyclins, including CycA, during cell entry into endocycles (***Sigrist and Lehner, 1997***). Remarkably, nearly all egg chambers in *bam>Ras^G12V^+uev1a-RNAi+fzr-RNAi* ovaries exhibited degradation (***Figure 5A, B***). Additionally, we knocked down three additional APC/C complex genes in *bam>Ras^G12V^+uev1a-RNAi* ovaries: *shattered* (*shtd*, encoding *Drosophila* APC1) **(*Tanaka-Matakatsu et al., 2007***), *morula* (*mr*, encoding *Drosophila* APC2) **(*Kashevsky et al., 2002***), and *lemming A* (*lmgA*, encoding *Drosophila* APC11) (***Nagy et al., 2012***). Among these factors, Shtd is a critical component of the scaffolding platform, while Mr and LmgA are essential components of the catalytic modules within the APC/C complex (***Yamano, 2019***). However, no significant enhancement in nurse cell death was observed, which may be due to the relatively mild phenotype of nurse cell death in *bam>Ras^G12V^+uev1a-RNAi* ovaries. Therefore, we switched to knocking down these genes in *nos>Ras^G12V^*ovaries. Notably, knockdown of any of them could significantly exacerbated *Ras^G12V^*-induced nurse cell death (***Figure 5C, D***), similar to the effect observed with Uev1A downregulation. Collectively, these findings provide genetic evidence that Uev1A collaborates with the APC/C complex to mitigate *Ras^G12V^*-induced nurse cell death.

**Figure 5.**
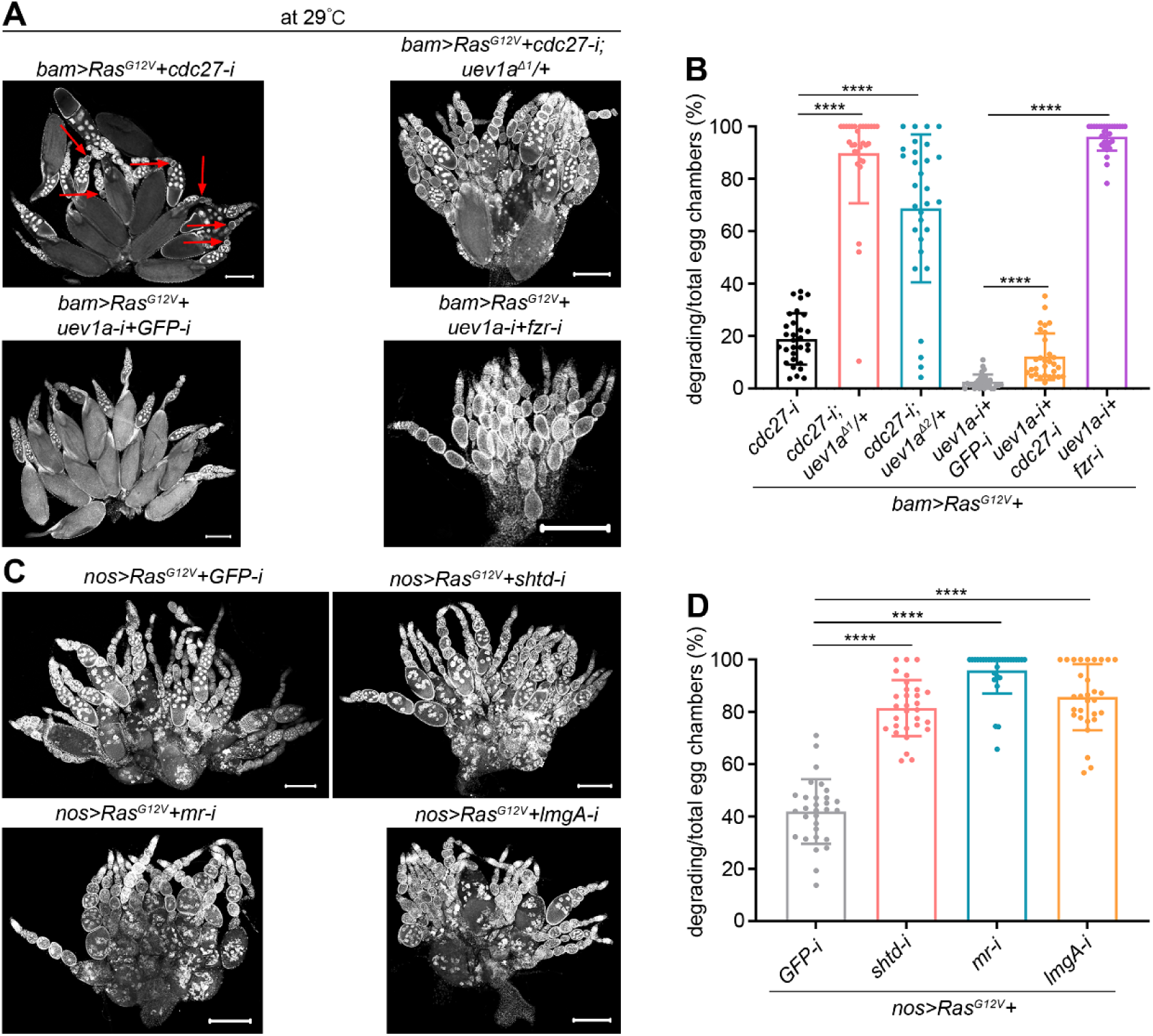
Uev1A collaborates with the APC/C complex to mitigate *Ras^G12V^*-induced nurse cell death. (A and C) Representative ovaries (DAPI staining). The red arrows in (A) denote degrading egg chambers. Scale bars: 200 μm. (B and D) Quantification data. 30 ovaries from 7-day-old (B) or 3-day-old (D) flies were quantified for each genotype. Statistical significance was determined using one-way ANOVA: **** (*P* < 0.0001). The online version of this article includes the following source data and figure supplements for figure 5: **Source data 2.** All genotypes. **Source data 3.** Raw quantification data. **Figure supplement 1.** Expression pattern of Uev1A in germarium. **Figure supplement 2.** Uev1A, Ben, and Cdc27 work together to protect nurse cells from death during normal oogenesis.

Then, we investigated whether Uev1A and the APC/C complex protect nurse cells from death during normal oogenesis. Given that the G2/M-promoting CycA is absent in the G/S-endocycling nurse cells (***Figure 4E***) (***Lilly et al., 2000; Lilly and Spradling, 1996***), we used the cystocyte-specific driver, *bam-GAL4-VP16*, to conduct the assays at 29°C. Similar to nurse cells, Uev1A expression was at basal levels in cystocytes, with some expression detected in inner sheath cells and epithelial follicle cells (***Figure 5—figure supplement 1***). Remarkably, nearly all egg chambers in *bam>fzr-RNAi* ovaries exhibited degradation (***Figure 5—figure supplement 2***), underscoring Fzr’s critical regulatory role in this context. Uev1A typically partners with Bendless (Ben), the *Drosophila* homolog of mammalian Ubc13 **(*Muralidhar and Thomas, 1993; Oh et al., 1994***), to perform the E2 enzyme function (***Sancho et al., 1998; Zhou et al., 2005***). Ovaries with the *bam>cdc27-RNAi*, *bam>cdc27-RNAi+uev1a-RNAi*, or *bam>cdc27-RNAi+ben-RNAi* genotype showed minimal degradation of egg chambers (***Figure 5—figure supplement 2***). However, degraded egg chambers were prevalent in *bam>cdc27-RNAi+ben-RNAi; uev1a*^Δ*1*^*/+* and *bam>cdc27-RNAi+ben-RNAi; uev1a*^Δ*2*^*/+* ovaries, where Uev1A, Ben, and Cdc27 are all downregulated (***Figure 5—figure supplement 2***). These results suggest that Uev1A, Ben, and the APC/C complex also work together to protect nurse cells from death during normal oogenesis.

### Uev1A and the APC/C complex work together to degrade CycA via the proteasome

The APC/C complex is known to collaborate with the E2 enzymes UBE2C (*Drosophila* homolog: Vihar) and UBE2S (*Drosophila* homolog: Ube2S) to facilitate the proteasomal degradation of CycA during cell cycle progression (***Greil et al., 2022; Yamano, 2019***). This degradation involves the assembly of branched ubiquitin chains, incorporating K11, K48, and K63 linkages through a two-step mechanism (***Meyer and Rape, 2014***). However, the involvement of Uev1A in this process remains unexplored. To explore it, we performed biochemical assays in cultured *Drosophila* Schneider 2 (S2) cells, overexpressing tag-fused proteins using *act-GAL4* to enhance expression efficiency. Co-immunoprecipitation (co-IP) assays revealed that Uev1A interacts with several components of the APC/C complex, including Mr (*Drosophila* APC2), Cdc16 (*Drosophila* APC6), and Cdc23 (*Drosophila* APC8). In addition, Cdc27 was found to interact with CycA (***Figure 6A, B and Figure 6—figure supplement 1***). Remarkably, RNAi targeting *uev1a*, *ben*, or *cdc27* significantly stabilized CycA proteins after the treatment with cycloheximide (CHX), an inhibitor of protein synthesis (***Figure 6C, D and Figure 6—figure supplement 2***). Since the K63 linkage primarily mediates lysosomal degradation via autophagy (***Kwon and Ciechanover, 2017***), we tested CycA stabilization with a chloroquine (CQ, a lysosome inhibitor) or MG132 (a proteasome inhibitor) treatment. Notably, MG132, but not CQ, treatment significantly stabilized CycA proteins (***Figure 6E***). These results suggest that Uev1A, Ben, and Cdc27 cooperate to promote CycA degradation through the proteasome rather than the lysosome.

**Figure 6.**
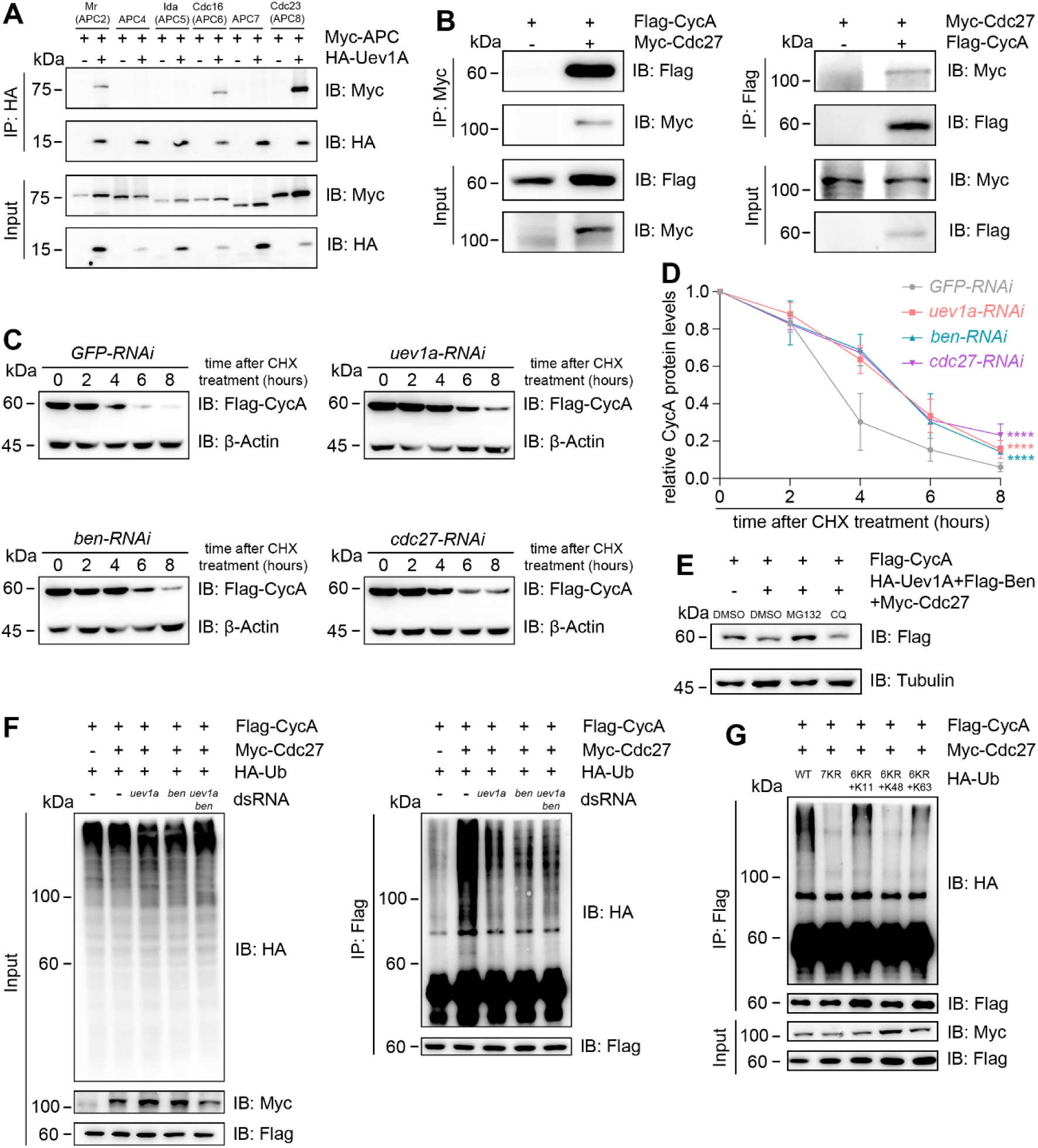
Uev1A, Ben, and Cdc27 work together to degrade CycA through the proteasome. (A and B) co-IP assays. The tagged proteins were co-expressed in S2 cells to assess physical interactions. As shown in (A), Uev1A interacts specifically with three APC/C subunits: Mr (APC2), Cdc16 (APC6), and Cdc23 (APC8). Assays in (B) demonstrate a physical interaction between CycA and Cdc27 (APC3). (C-E) CycA stability assays. CHX: a protein-synthesis inhibitor; MG132: a proteasome inhibitor; CQ: a lysosome inhibitor. In (D), the relative levels of CycA proteins were quantified using the following formula: (Mean gray value of the CycA/β-Actin band at n hours post-treatment) ÷ (Mean gray value of the CycA/β-Actin band at 0 hour). Three independent replicates were conducted at each timepoint, and statistical significance was determined using two-way ANOVA with multiple comparisons: **** (*P* < 0.0001). (F and G) CycA ubiquitination assays in S2 cells. As shown in (F), Cdc27 promotes CycA ubiquitination in a Uev1A/Ben-dependent manner. Assays in (G) indicate that the K11 and K63 of ubiquitin are required for CycA ubiquitination. The online version of this article includes the following source data and figure supplement for figure 6: **Source data 3.** Raw quantification data. **Figure supplement 1.** Co-IP results. **Figure supplement 2.** RNAi efficiency assays.

To directly validate this, we performed ubiquitination assays for CycA in S2 cells. Cdc27 significantly enhanced CycA ubiquitination, while this effect was markedly reduced upon the knockdown of *uev1a*, *ben*, or both (***Figure 6F***). This result indicates that both Uev1A and Ben are essential for APC/C-mediated proteasomal degradation of CycA. Furthermore, we explored the roles of the seven lysine (K) residues in ubiquitin for polyubiquitin chain formation. The 7KR (lysine to arginine) mutation completely abolished Cdc27-promoted polyubiquitination of CycA. Intriguingly, the 6KR+K48 mutation, in which all K residues except K48 were mutated to R residues, failed to restore polyubiquitination. In contrast, either 6KR+K11 or 6KR+K63 mutation significantly restored polyubiquitination (***Figure 6G***). These findings suggest that the K11 and K63 linkages are primarily responsible for CycA polyubiquitination in *Drosophila* cells. Together with previous studies (***Hofmann and Pickart, 1999; McKenna et al., 2001***), we propose that Uev1A and Ben mediate the K63 linkage in this process.

### Uev1A inhibits the overgrowth of *Drosophila* germline tumors driven by oncogenic *Ras^G12V^*

Given the absence of cell division in normal polyploid nurse cells **(*Hammond and Laird, 1985***), their death induced by division-promoting *Ras^G12V^* represents an artificial stress. Notably, Uev1A was not upregulated in response to this stress (***Figure 3E***), and it executed the function through degrading CycA (***Figure 4-6***). These findings prompted us to investigate whether Uev1A also counteracts oncogenic *Ras*-driven tumorigenesis in diploid cells, which undergo normal cell division. Our prior research demonstrated that oncogenic *Ras^G12V^*markedly promotes the overgrowth of diploid *bam*-deficient germline tumors (***Zhang et al., 2024***), which are highly resistant to cell death (***Zhang et al., 2023; Zhao et al., 2018***). Intriguingly, knocking down *uev1a* significantly enhanced the overgrowth of these tumors, while overexpressing Uev1A suppressed it (***Figure 7A, B***).

**Figure 7.**
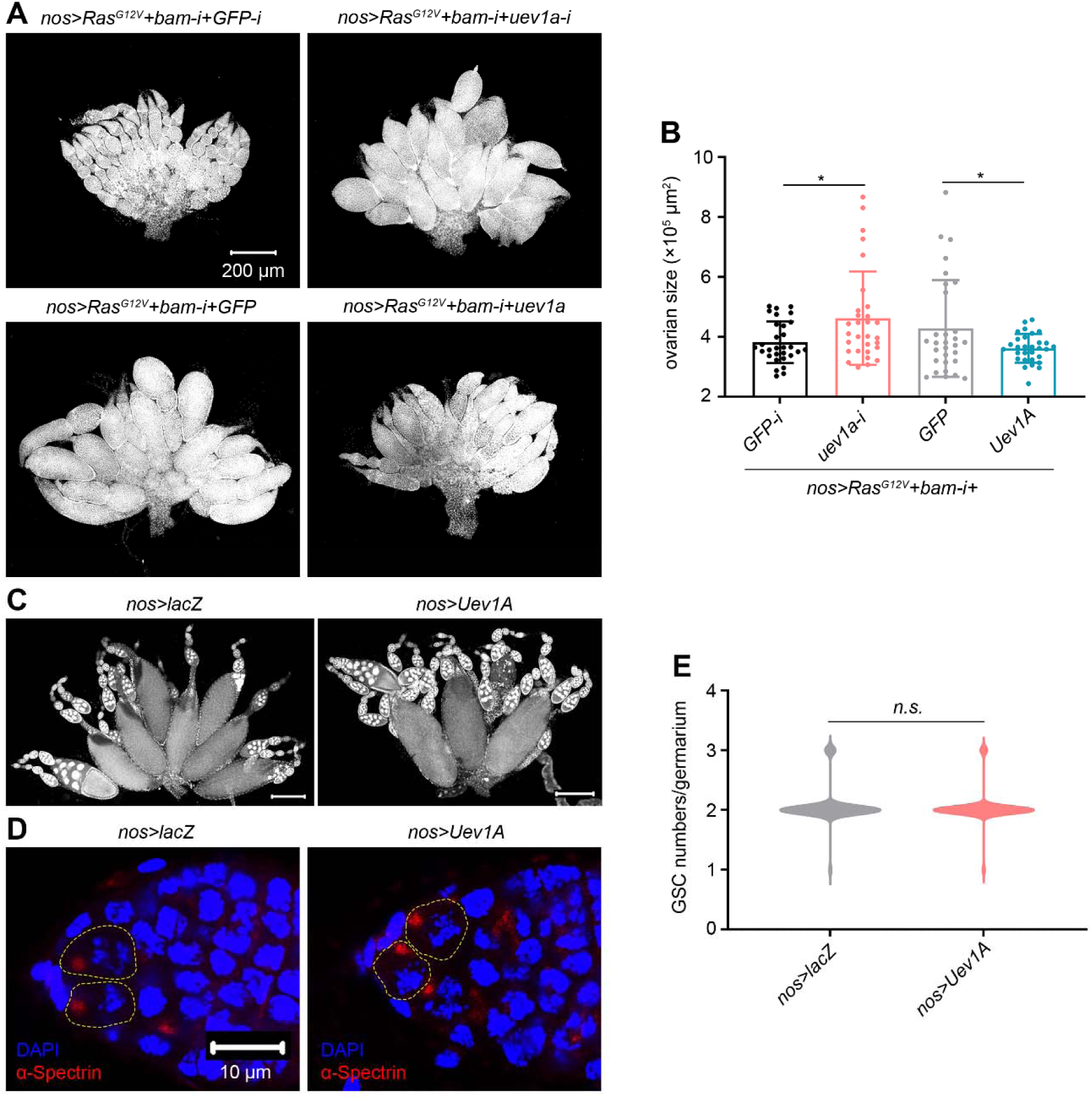
Uev1A inhibits the overgrowth of germline tumors induced by oncogenic *Ras^G12V^*. (A and C) Representative ovaries (DAPI staining). All images in (A) are of the same magnification. Scar bars in (C): 200 μm. (B) Quantification data for ovarian size. The largest 2D area of each ovary in a single confocal focal plane was scanned, and its size was measured using ImageJ. 30 ovaries from 3-day-old flies were analyzed for each genotype. (D) Representative samples. The GSCs within stem cell niches are outlined by yellow dashed lines. Both images are of the same magnification. (E) Quantification data for GSC numbers per germarium. Germ cells that directly contact cap cells and contain dot-like spectrosomes were counted as GSCs. 100 germaria from 14-day-old flies were quantified for each genotype. In (B and E), statistical significance was determined using t test: *n.s.* (*P* > 0.05) and * (*P* < 0.05). The online version of this article includes the following source data for figure 7: **Source data 2.** All genotypes. **Source data 3.** Raw quantification data.

These results indicate that Uev1A also plays a role in counteracting oncogenic *Ras*-driven tumorigenesis in diploid cells.

Considering the therapeutic potential of gene upregulation in inhibiting tumor growth, a critical concern is its impact on normal physiological processes. In this study, we examined the impact of Uev1A overexpression on *Drosophila* oogenesis and GSC maintenance. Notably, the *nos>Uev1A* flies remained fertile, and their ovaries appeared morphologically similar to those of the *nos>lacZ* control flies (***Figure 7C***), indicating normal oogenesis. Furthermore, each germarium in the ovaries of 14-day-old *nos>Uev1A* flies contained a similar number of GSCs as those in the *nos>lacZ* control flies (***Figure 7D, E***). These results suggest that Uev1A overexpression does not disrupt normal oogenesis and GSC maintenance.

### UBE2V1 and UBE2V2 inhibit the overgrowth of human colorectal tumors driven by oncogenic *KRAS*

Our findings in *Drosophila* prompted us to explore the tumor-suppressive effects of UBE2V1 and UBE2V2 on the growth of *RAS*-mutant human tumors. Using the Kaplan-Meier plotter (https://kmplot.com/analysis), we first evaluated the correlation between UBE2V1/2 expression and prognosis in several types of *RAS*-mutant cancer patients, including melanoma, myeloma, lung cancer, and colorectal cancer. Among them, higher expression levels of UBE2V1/2 were significantly associated with improved relapse-free survival in *KRAS*-mutant colorectal cancer patients (***Figure 8A***). RNA-seq data from The Cancer Genome Atlas (TCGA) showed that *UBE2V1* and *UBE2V2* are transcribed at similar levels in colorectal cancer patients with oncogenic *RAS* mutations (including *KRAS*, *HRAS*, and *NRAS*) as in those without such mutations (***Figure 8—figure supplement 1***). These findings suggest that UBE2V1 and UBE2V2 are not upregulated in response to oncogenic *RAS* mutations, paralleling the behavior of Uev1A in *Drosophila* ovarian nurse cells (***Figure 3E***).

**Figure 8.**
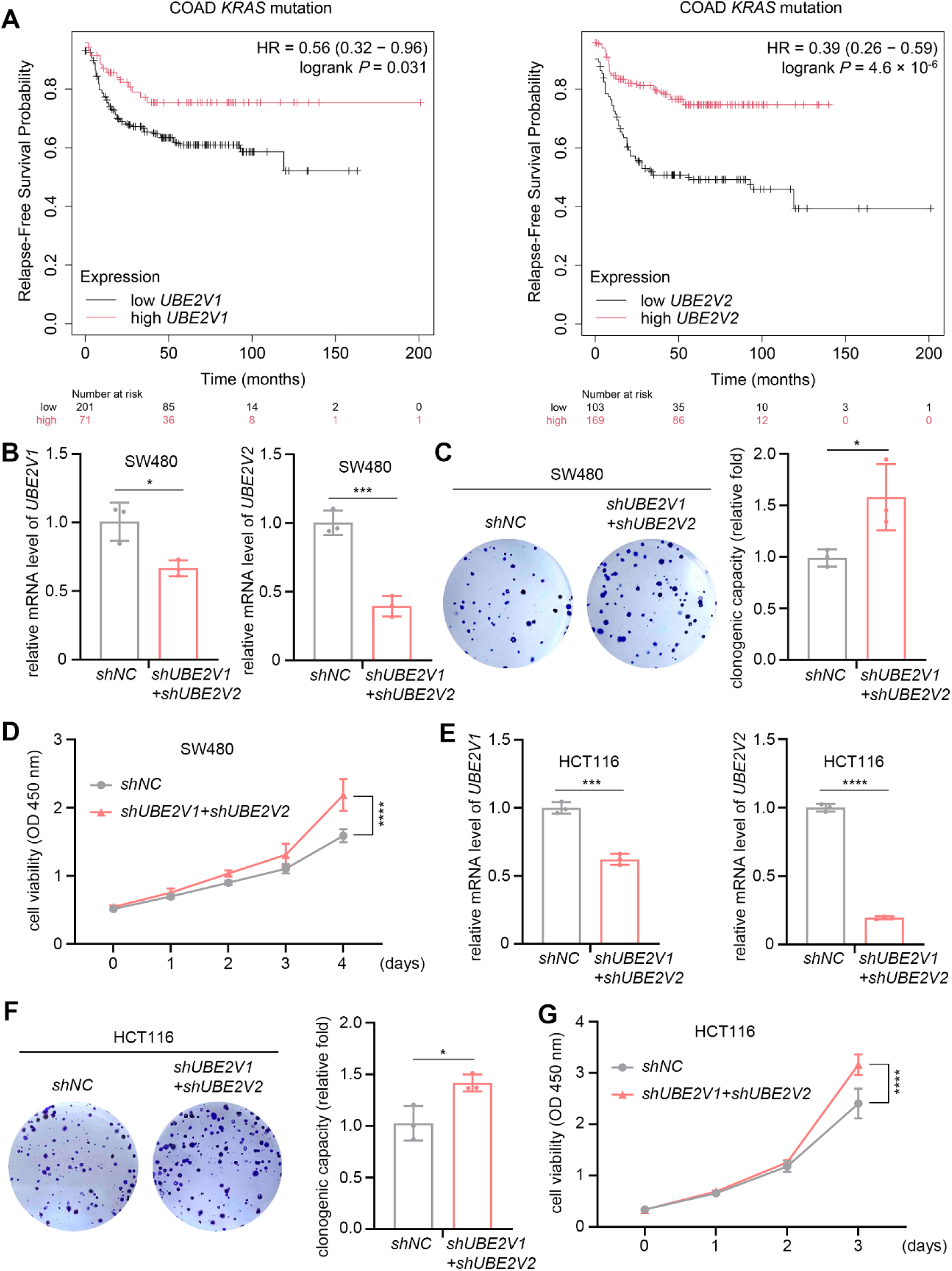
Prognostic significance and tumor-suppressive effects of UBE2V1 and UBE2V2 on *KRAS*-mutant colorectal cancer. (A) Kaplan-Meier analysis of relapsefree survival in *KRAS*-mutant colorectal cancer patients with high or low expression levels of UBE2V1 and UBE2V2. (B and E) The knock-down efficiency assays. The relative mRNA levels were normalized to *GAPDH*. (C-G) Assays to evaluate the effects of *UBE2V1-* and *UBE2V2-RNAi* on colony formation and cell viability in SW480 and HCT116 cells. In (B, C, E, and F), three independent replicates were conducted, and statistical significance was determined using t test. In (D and G), five (D) and six (G) independent replicates were conducted at each time point, and statistical significance was determined using two-way ANOVA with multiple comparisons. * (*P* < 0.05), *** (*P* < 0.001), and **** (*P* < 0.0001). The online version of this article includes the following source data and figure supplement for figure 8: **Source data 3.** Raw quantification data. **Figure supplement 1.** TCGA analysis comparing UBE2V1 and UBE2V2 expression levels in colorectal cancer patients with and without *RAS* mutations. **Figure supplement 2.** Knocking down either *UBE2V1* or *UBE2V2* alone mildly influences the growth of colorectal cancer cell lines.

To investigate the potential tumor-suppressive roles of UBE2V1 and UBE2V2 in *KRAS*-mutant colorectal cancer, we performed individual knockdowns of each gene in two human colorectal cancer cell lines: SW480 cells (carrying the *KRAS^G12V^* mutation) and HCT116 cells (carrying the *KRAS^G13D^* mutation). Notably, individual knockdown of either *UBE2V1* or *UBE2V2* only mildly affected the proliferation of SW480 cells (***Figure 8-figure supplement 2***), suggesting potential functional redundancy between the two proteins. However, combined knockdown of *UBE2V1* and *UBE2V2* significantly enhanced colony formation and cell viability in both SW480 and HCT116 cells, as demonstrated by clonogenic and CCK8 assays (***Figure 8B-G***). These results indicate that UBE2V1 and UBE2V2 exert tumor-suppressive effects in colorectal cancer cells harboring oncogenic *KRAS* mutations.

Next, we examined whether overexpression of UBE2V1 or UBE2V2 could suppress the growth of SW480 and HCT116 cells. The overexpression efficiency of each protein was confirmed by Western blotting (***Figure 9—figure supplement 1A***). Using 5-ethynyl-2’-deoxyuridine (EdU) incorporation assays, we found that overexpression of either UBE2V1 or UBE2V2 significantly inhibited the proliferation of both cancer cell lines *in vitro* (***Figure 9—figure supplement 2A, B***). This antiproliferative effect was further supported by clonogenic and cell viability assays (***Figure 9—figure supplement 2C-E***).

**Figure 9.**
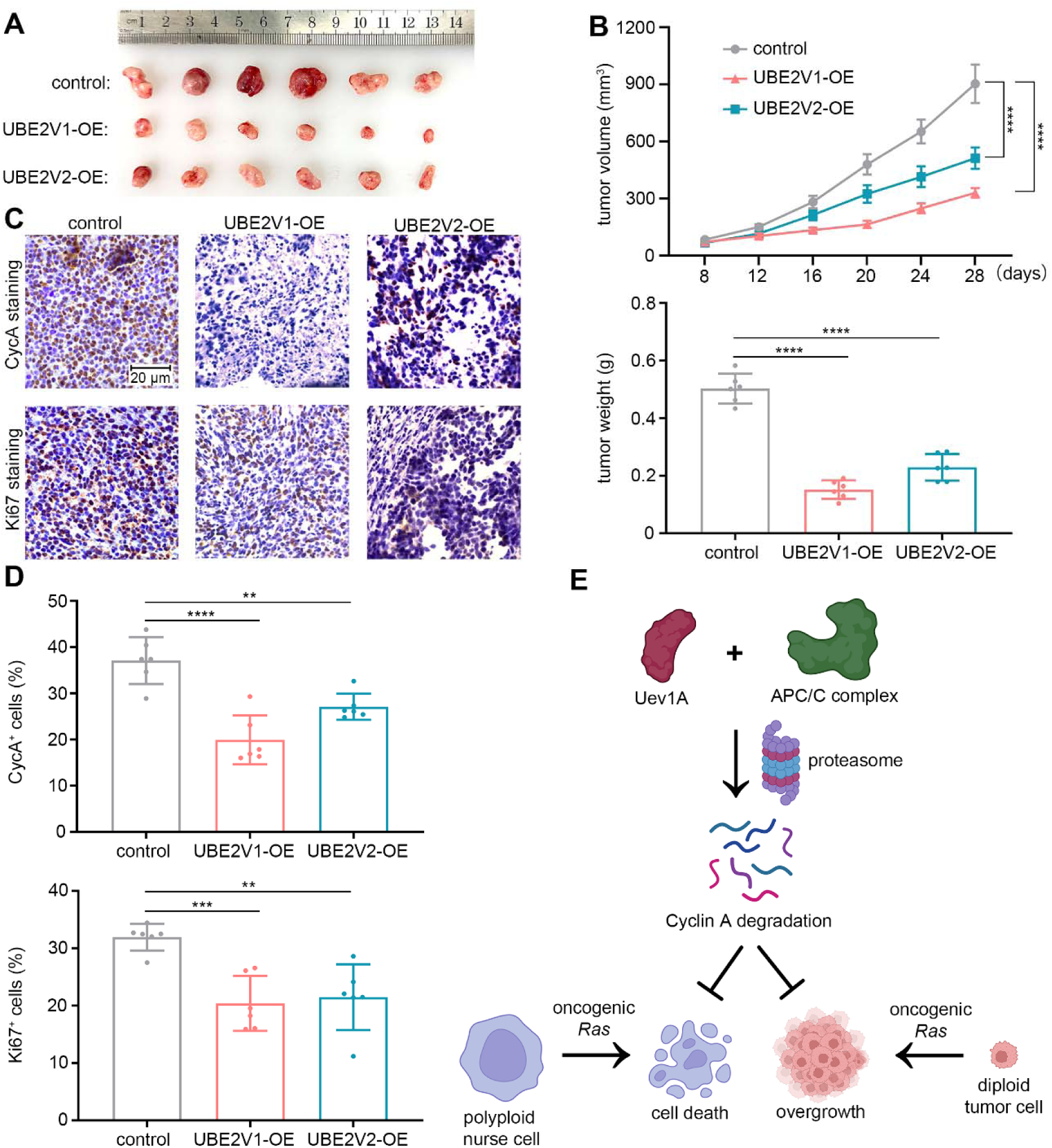
Overexpression of UBE2V1 or UBE2V2 suppresses the growth of *KRAS*-mutant colorectal cancer. (A and B) Subcutaneous tumorigenesis assays in nude mice, where tumors were excised, photographed, and weighed 28 days after tumor cell injection. (C and D) Immunohistochemical staining to assess CycA expression and Ki-67 positivity in tumor tissues. All images in (C) are of the same magnification. In (B-1), six independent replicates were conducted, and statistical significance was determined using two-way ANOVA with multiple comparisons. In (B-2 and D), six independent replicates were conducted, and statistical significance was determined using one-way ANOVA. ** (*P* < 0.01), *** (*P* < 0.001), and **** (*P* < 0.0001). (E) Working model. By degrading CycA, Uev1A and the E3 APC/C complex counteract oncogenic *Ras* stimuli, thereby protecting against cell death in polyploid *Drosophila* nurse cells and suppressing overgrowth in diploid *Drosophila* germline and human colorectal tumor cells. The online version of this article includes the following source data and figure supplement for figure 9: **Source data 3.** Raw quantification data. **Figure supplement 1.** Validation of UBE2V1/2 overexpression in colorectal cancer cell lines. **Figure supplement 2.** UBE2V1/2 overexpression suppresses the growth of colorectal cancer cell lines.

To validate the tumor-suppressive effects of UBE2V1 and UBE2V2 *in vivo*, we established stable SW480 cell lines overexpressing each protein, with overexpression confirmed by Western blotting (***Figure 9—figure supplement 1B***). In subcutaneous tumorigenesis assays using *Balb/c* nude mice, overexpression of either UBE2V1 or UBE2V2 significantly inhibited tumor formation compared to control (***Figure 9A, B***). Immunohistochemical analysis of the resulting tumors showed a marked decrease in Cyclin A expression and Ki-67-positive cells, indicating reduced proliferation (***Figure 9C, D***). Notably, the tumor-suppressive effect of UBE2V1 was more robust than that of UBE2V2, consistent with the stronger protective effect of UBE2V1 against *Ras^G12V^*-induced nurse cell death in *Drosophila* (***Figure 2F, G***). Taken together, these findings provide strong evidence for the tumor-suppressive roles of UBE2V1 and UBE2V2 in *KRAS*-mutant colorectal cancer.

## Discussion

It is well established that oncogenic *Ras* can induce DNA replication stress, leading to cellular senescence or death (***Hills and Diffley, 2014; Kotsantis et al., 2018***). However, our previous (***Zhang et al., 2024***) and current findings demonstrated that oncogenic *Ras^G12V^* can also trigger cell death in polyploid *Drosophila* ovarian nurse cells through aberrantly promoting their division. In this study, we performed a genome-wide genetic screen and identified the E2 enzyme Uev1A as a crucial protector against this specific form of cellular stress. Mechanistically, Uev1A collaborates with the APC/C complex (E3) to facilitate the proteasomal degradation of CycA, which overexpression alone can also trigger nurse cell death. Furthermore, Uev1A and its human homologs, UBE2V1 and UBE2V2, counteract oncogenic *Ras*-driven tumorigenesis in diploid *Drosophila* germline and human colorectal tumor cells, respectively. These findings highlight the critical role of Uev1A in counteracting oncogenic *Ras* stimuli in both polyploid and diploid cells (***Figure 9E***).

Our studies reveal that the DDR pathway protects against *Ras^G12V^*-induced nurse cell death (***Figure 3A, B***), suggesting that this cellular stress both activates and is subsequently suppressed by the DDR. In contrast, p53 promotes nurse cell death under the same conditions (***Figure 3A, B***). Interestingly, previous research has also demonstrated distinct roles of p53 and Lok, a crucial DDR regulator, in regulating nurse cell death during mid-oogenesis. While Lok overexpression triggers nurse cell death, p53 overexpression does not. Furthermore, Lok-induced nurse cell death remains unaffected by *p53* mutation, indicating that its mechanism operates independently of p53 (***Bakhrat et al., 2010***). The underlying mechanisms driving these differences warrant further investigation.

Of note, the suppressive effects of Uev1A on *Ras^G12V^; bam^RNAi^* germline tumors in *Drosophila* were less pronounced than that of UBE2V1/2 on oncogenic *KRAS^G12V^*-driven human colorectal tumor xenografts in nude mice (***compare Figure 7A, B with Figure 9A, B***). Our previous research has shown that *Ras^G12V^; bam^-/-^* germline tumor cells divide infrequently (***Zhang et al., 2024***). This suggests that the relatively weak suppression observed in these tumors may be due to the fact that Uev1A’s mechanism of action is cell cycle-dependent: the more frequently tumor cells divide, the more effectively Uev1A can suppress tumor overgrowth. We did not observe significant negative effects of Uev1A overexpression on GSC maintenance in *Drosophila* ovaries (***Figure 7D, E***). This is likely because GSCs also divide infrequently (***Morris and Spradling, 2011***). Therefore, these findings underscore the importance of considering the potential negative effects of Uev1A/UBE2V1/2 overexpression in other contexts, particularly when cells divide rapidly.

RAS oncoproteins have long been considered “undruggable”, due to the lack of deep-binding, targetable pockets (***Moore et al., 2020***). To date, targeted therapies have been approved only for KRAS^G12C^ **(*Canon et al., 2019; Ou et al., 2022; Skoulidis et al., 2021***). Recently, small-molecule pan-KRAS degraders, developed using the proteolysis-targeting chimeras (PROTACs) strategy, have shown promise (***Popow et al., 2024***), though their clinical effectiveness and safety remain to be fully validated. Alternatively, targeting key regulators in the RAS signaling pathway—such as SOS, SHP2, Farnesyltransferase, Raf, MEK, ERK, and PI3K—has emerged as an attractive therapeutic approach (***Moore et al., 2020***). Recent research has also explored the strategy of hyperactivating oncogenic *RAS* to trigger cell death as a potential means to suppress the overgrowth of human colon tumor xenografts in nude mice (***Dias et al., 2024***). Additionally, targeting the cellular dependencies driven by oncogenic *RAS* may trigger RAS-specific synthetic lethality, presenting a promising therapeutic approach **(*Moore et al., 2020***). Our findings highlight the tumor-suppressive effects of UBE2V1 and UBE2V2 on human colorectal tumors driven by oncogenic *KRAS* (***Figure 8 and 9***). Notably, both of these two E2 enzymes are relatively small, with UBE2V1 consisting of 147 amino acid residues and UBE2V2 145 residues (***Figure 2E***). This raises the exciting possibility that their upregulation, through mRNA delivery or small-molecule agonists, could offer a promising therapeutic approach for human cancer.

## Materials and Methods

### Key resources table

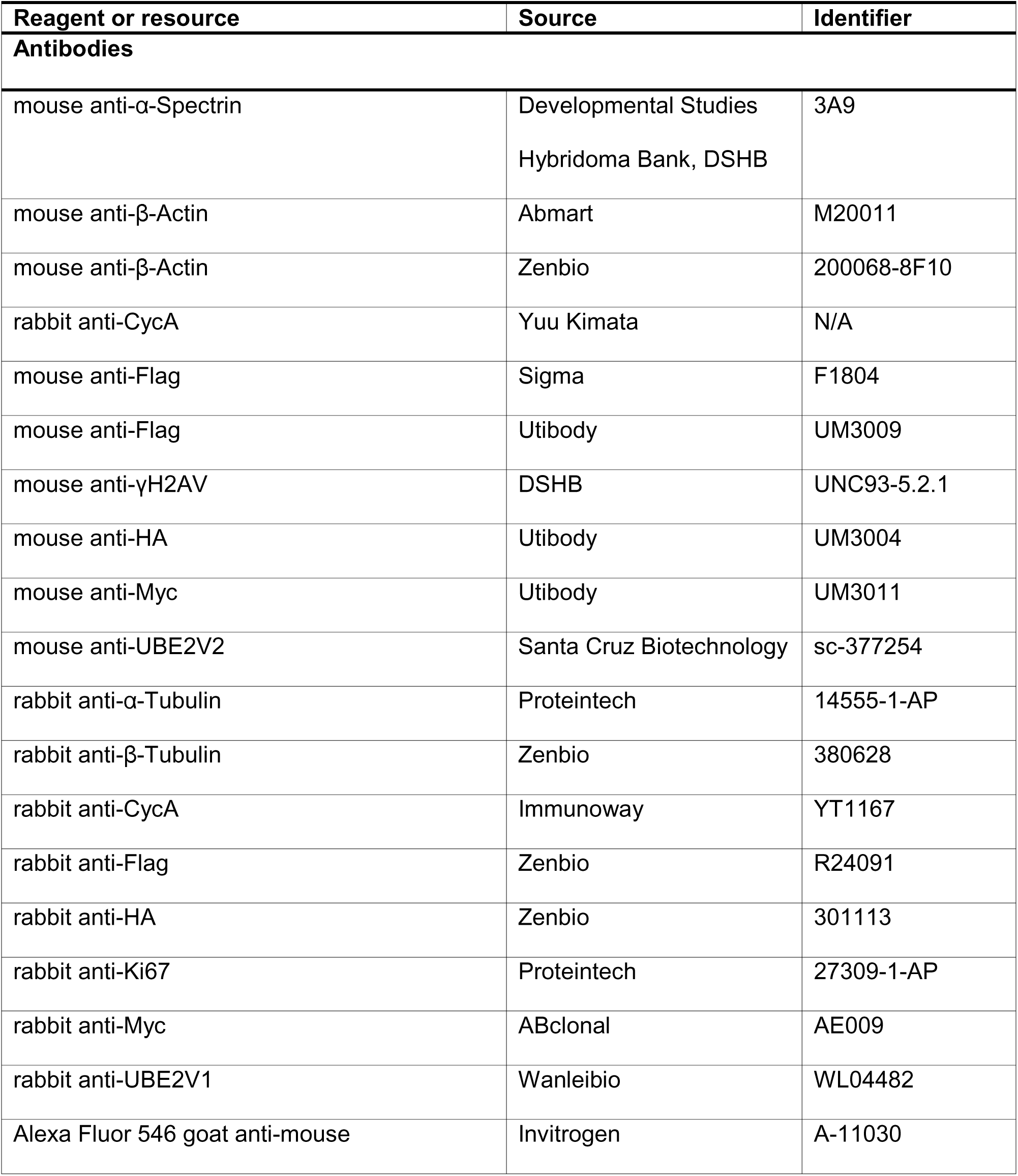
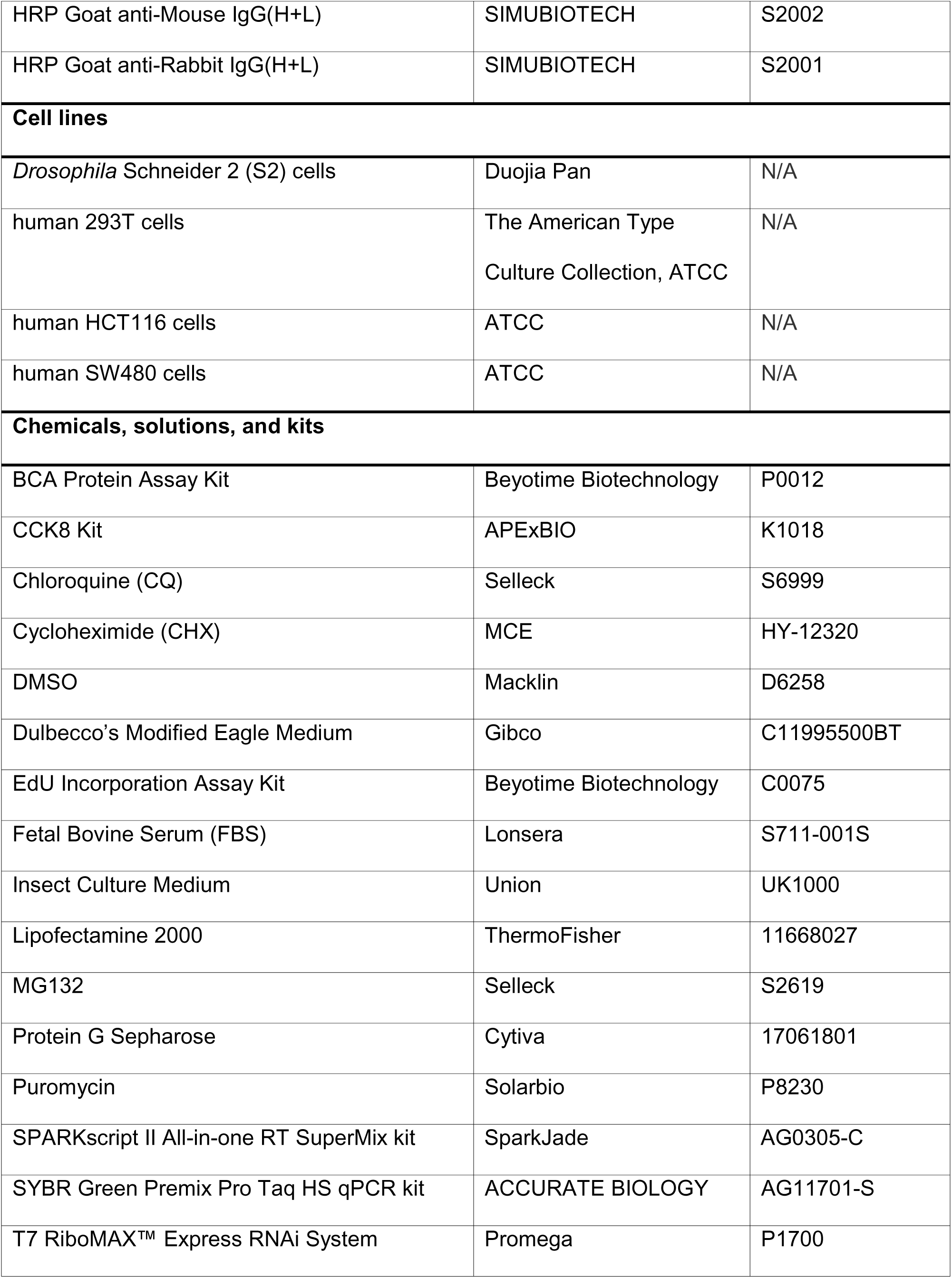
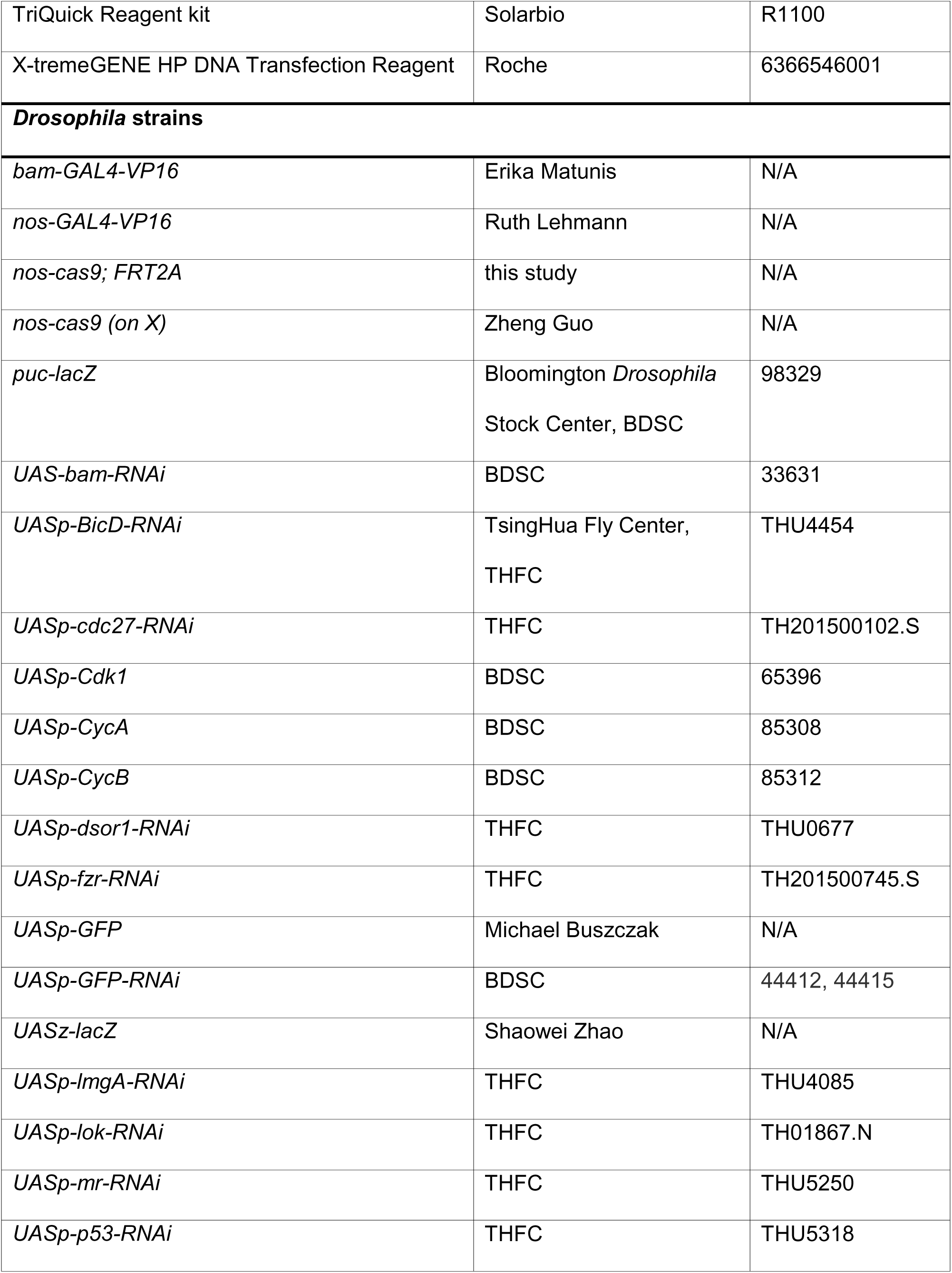
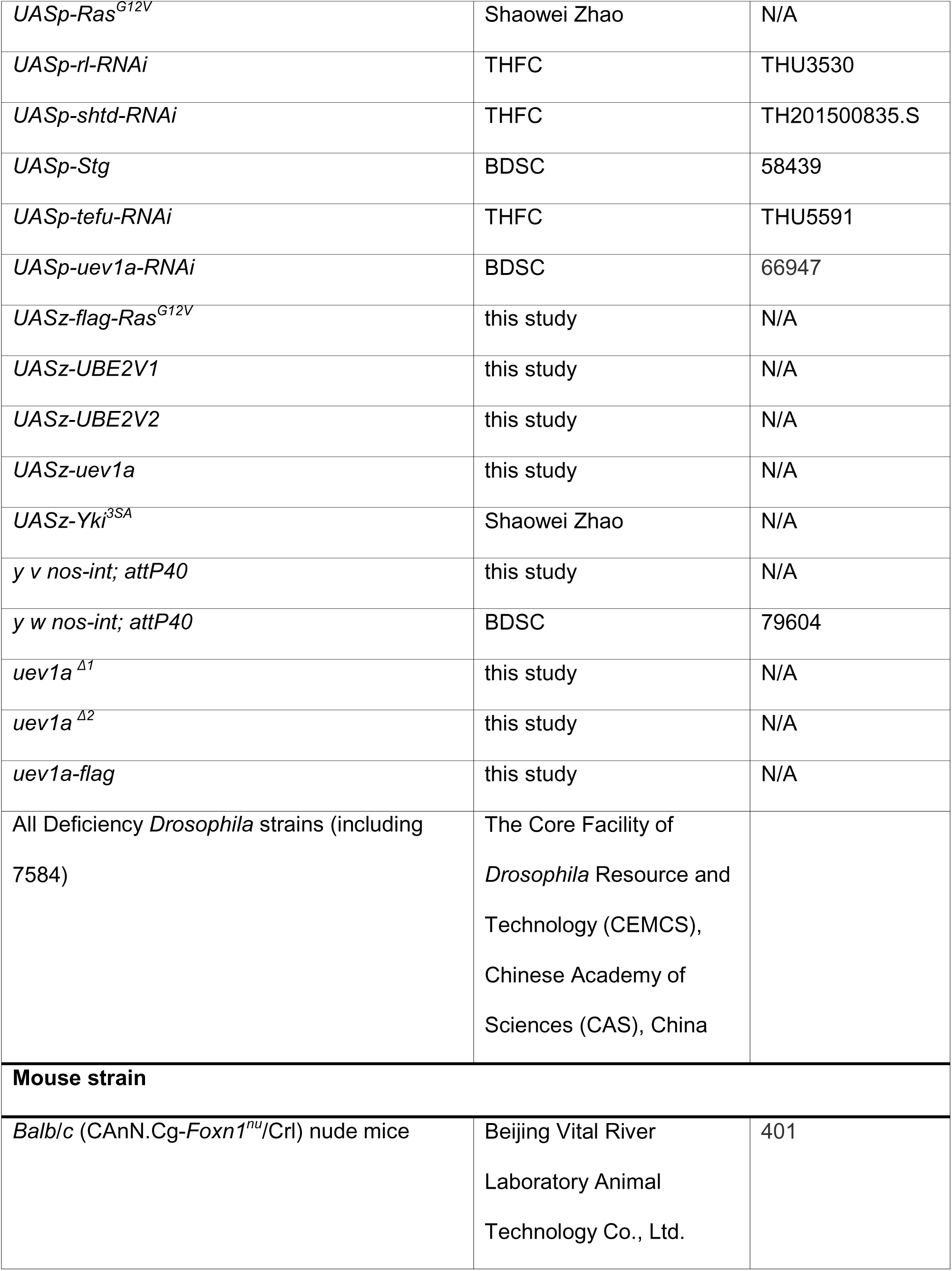
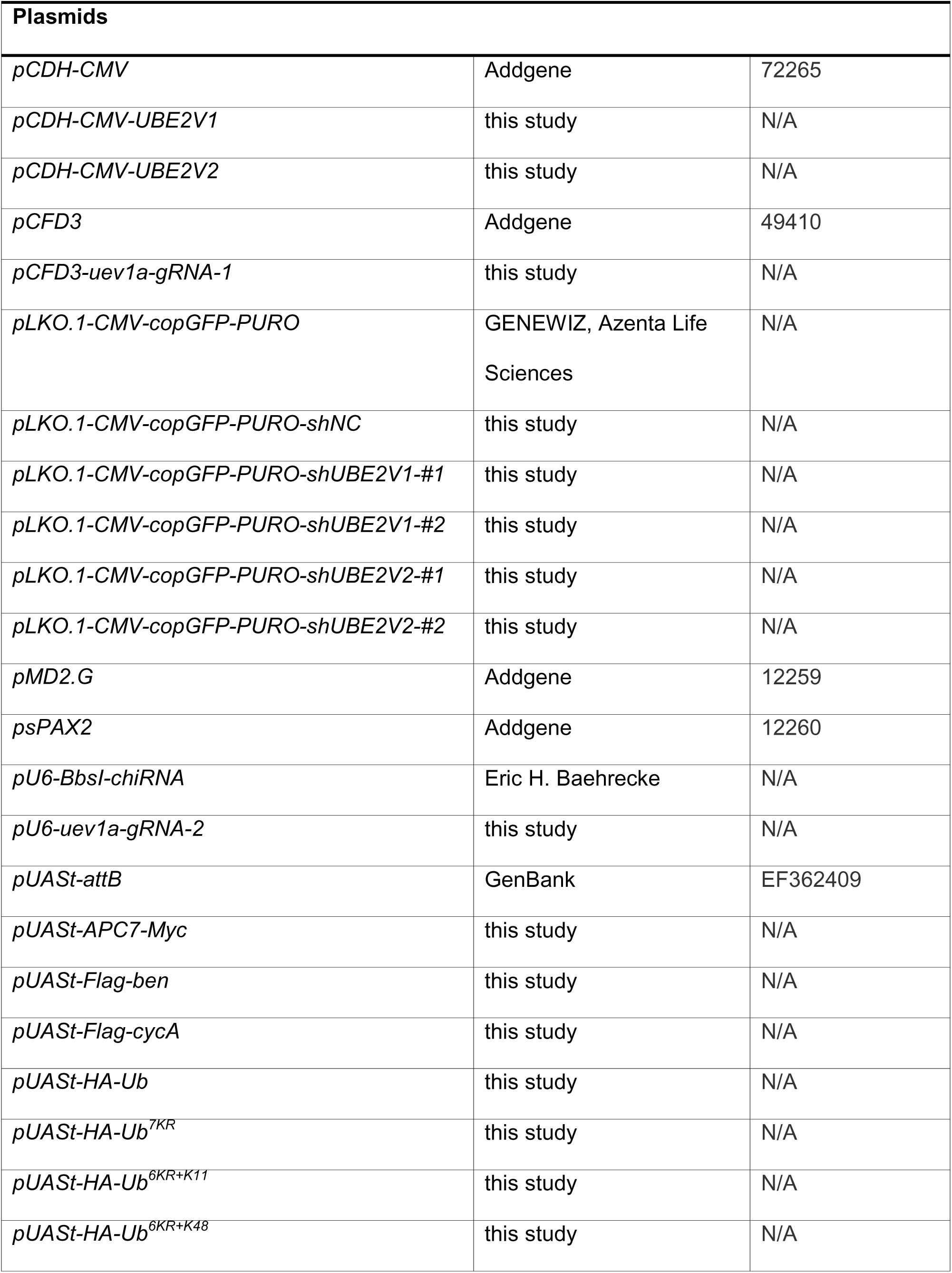
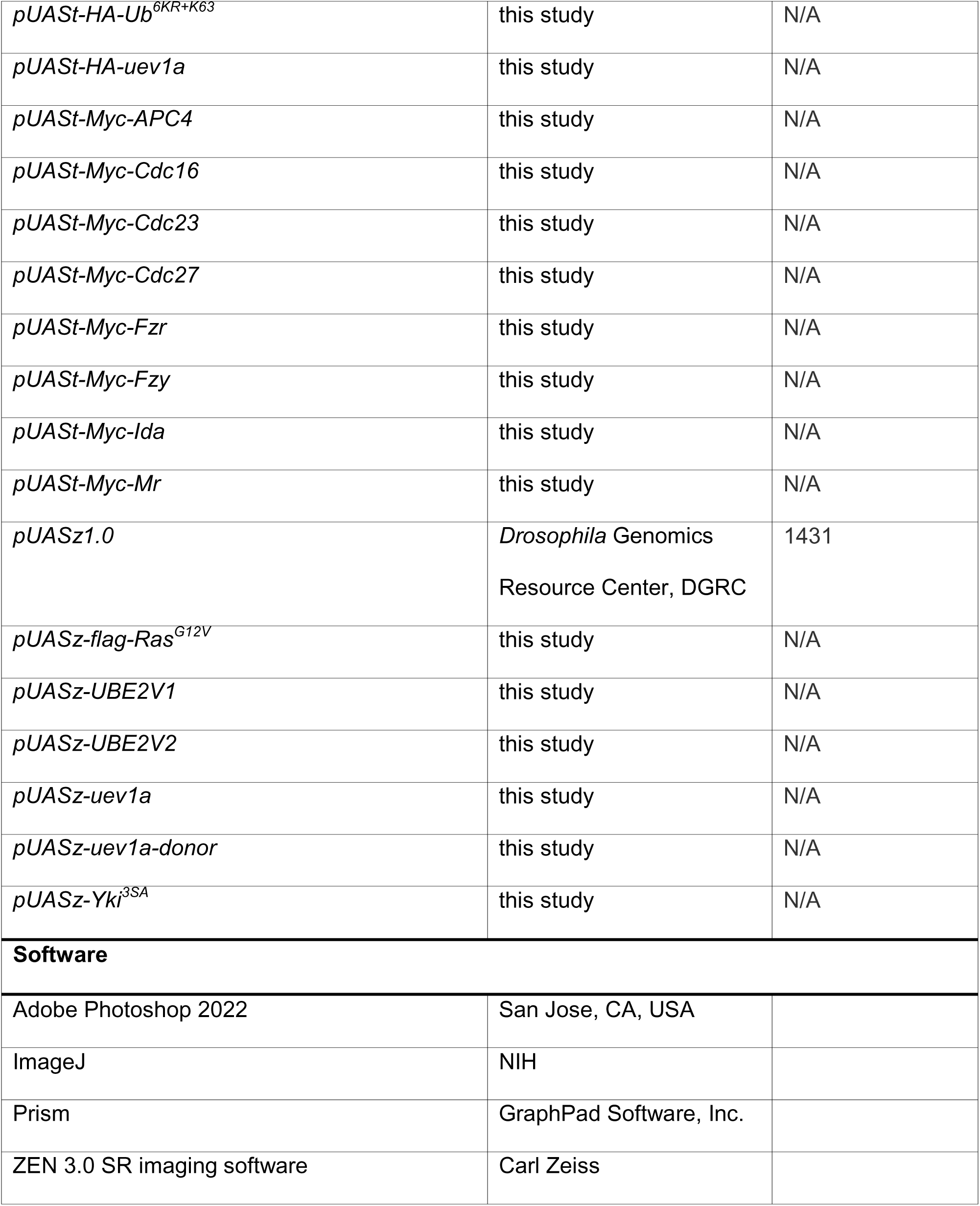

## Data availability

The results of the deficiency screening are provided in **Source data 1**. All genotypes are listed in **Source data 2**, and the raw quantification data can be found in **Source data 3**.

### Fly husbandry

Cross experiments were conducted at 25°C, except for some performed at 29°C as noted.

### Transgenic flies

#### UASz-flag-Ras^G12V^, UASz-uev1a, UASz-UBE2V1, and UASz-UBE2V2

The coding sequences (CDSs) of *flag-Ras^G12V^*, *uev1a-RA*, *UBE2V1*, and *UBE2V2* were cloned into the *pUASz1.0* vector (***DeLuca and Spradling, 2018***). These plasmids were then microinjected into fertilized fly embryos with the genotype “*y w nos-int; attP40*”, resulting in the generation of transgenic fly strains. The DNA sequence encoding 3xFlag was as follows: ATGGACTACAAAGACCATGACGGTGATTATAAAGATCATGACATCGATTACAAGGATGACGATGACAAGCTT.

#### uev1a ^Δ1^ and uev1a ^Δ2^ mutants

The gRNA targeting the second coding exon of the *uev1a* gene was designed with the DNA sequence “GATCCAGCTCCTCCAGTAAGCGG” and cloned into the *pCFD3* vector (***Port et al., 2014***). The plasmids were then microinjected into fertilized fly embryos with the genotype “*y v nos-int; attP40*” to generate transgenic fly strain. These transgenic flies were subsequently crossed with “*nos-cas9; FRT2A*” flies to generate *uev1a* knockout mutants, which molecular information was identified by Sanger sequencing.

#### uev1a-flag knock in

The gRNA targeting the second-to-last coding exon of the *uev1a* gene was designed with the DNA sequence “GTTATAAACCGGAGCGTTGGTGG” and cloned into the *pU6-BbsI-chiRNA* vector. The homology arms consisted of 991 bp upstream of the stop codon (“TAG”), followed by a 3xFlag tag sequence, the “TAG” stop codon, and 979 bp downstream of “TAG”. To prevent cleavage by the gRNA, the targeting sequence within the upstream homology arm was mutated to “CCTACCCTGCGCTTCATTA”, ensuring that the amino acid sequence remained unaltered. These DNA sequences were synthesized into the *pUASz1.0* vector between the *BamHI* and *KpnI* sites and used as a double-stranded DNA (dsDNA) donor for homologous recombination-mediated repair. The *pU6-uev1a-gRNA-2* (100 ng/μL) and *pUASz-uev1a-donor* (100 ng/μL) plasmids were then co-injected into fertilized fly embryos with the genotype “*nos-cas9 (on X)*”. The resulting *uev1a-flag* knock-in fly strain was identified by PCR and confirmed by Sanger sequencing.

### Immuno-fluorescent staining

Fly ovaries were dissected in the PBS solution, fixed by the 4% paraformaldehyde solution (diluted in PBS) for 3 hours, washed by the PBST solution (0.3% Triton X-100 diluted in PBS) for 1 hour at room temperature (RT), then incubated with the primary antibodies (diluted in PBST) overnight at 4°C, washed by the PBST solution for 1 hour at RT, incubated with the secondary antibodies and 0.1 μg/mL DAPI (diluted in PBST) overnight at 4°C, washed by the PBST solution for 1 hour at RT, and subsequently mounted using 70% glycerol (autoclaved). The primary antibodies were used at the concentrations as follows: rabbit anti-CycA at 1:1000, mouse anti-Flag (Sigma, #F1804) at 1:500, mouse anti-γH2AV at 1:200, and mouse anti-α-Spectrin at 1:100. The Alexa Fluor conjugated secondary antibodies were used at 1:2,000.

### Construction of plasmids used in S2 cells

The CDSs of the genes were amplified from the cDNAs derived from either S2 cells or adult wild-type (*w^1118^*) flies and subsequently cloned into the *pUASt-attB* vector. N- or C-terminal tags were incorporated based on prior studies or structural predictions from AlphaFold. The following proteins were tagged at the N-terminus: APC4, Ben, CDC16, CDC23, Cdc27, CycA, Fzr, Fzy, Ida, Mr, Ub, Ub^7KR^, Ub^6KR+K11^, Ub^6KR+K48^, Ub^6KR+K63^, and Uev1A. In contrast, APC7 was tagged at the C-terminus.

### Culture and transfection of S2 cells

S2 cells were cultured in insect medium with 10% FBS and incubated at 27°C without CO₂. To transfect, cells were plated in 6-well plates to reach 70% confluence and incubated for 3 hours at 27°C. Then, a pre-complexed mixture of plasmid DNA and X-tremeGENE HP DNA transfection reagent was added slowly to the cultures. The transfection process followed the manufacturer’s instructions. After gentle mixing, cells were incubated at 27°C, and samples were collected 36 hours after transfection.

### Immunoprecipitation (IP), immunoblotting (IB), ubiquitination and protein stability assays in S2 cells

#### IP assay

The transfected S2 cells were lysed using NP-40 lysis buffer supplemented with protease inhibitors at 4°C for 30 minutes. The supernatants were incubated with primary antibodies at 4°C for 3 hours, followed by incubation with Protein G Sepharose for 2 hours. The beads were washed five times with NP-40 lysis buffer and boiled for 5 minutes in 2×SDS protein loading buffer [0.25 M Tris-HCl (pH 6.8), 78 mg/mL DTT, 100 mg/mL SDS, 50% glycerol, and 5 mg/mL bromophenol blue]. Then the samples were subjected to Western blotting. The following antibodies were used: mouse anti-Flag (Utibody, #UM3009) at 1:200, mouse anti-Myc at 1:200, and mouse anti-HA at 1:200.

#### IB assay

After 36 hours of transfection, the S2 cells were harvested and lysed in 2×SDS protein loading buffer. The lysates were then boiled for 5 minutes and subjected to Western blotting. The following antibodies were used: mouse anti-Flag (Utibody, #UM3009) at 1:5000, rabbit anti-Flag at 1:5000, rabbit anti-Myc at 1:3000, rabbit anti-HA at 1:3000, mouse anti-β-Actin (Zenbio, #200068-8F10) at 1:5000, and rabbit anti-β-Tubulin at 1:5000.

#### Ubiquitination assay

S2 cells were transiently transfected with the indicated plasmid combinations. 30 hours post-transfection, cells were treated with 50 μM MG132 for 6 hours. Proteins were then immunoprecipitated and analyzed by Western blotting.

#### Protein stability assay

S2 cells were treated with 20 µg/mL of the protein synthesis inhibitor CHX for specified time intervals prior to harvesting.

### RNAi assay in S2 cells

Two double-stranded RNAs (dsRNAs) targeting distinct regions of the relevant genes were synthesized using the T7 RiboMAX™ Express RNAi System. S2 cells were seeded in 6-well tissue culture plates and treated with 15 μg of dsRNA in the culture medium, followed by incubation for 24 hours. Expression plasmids were then transiently transfected into the cells, after which an additional 10 μg of dsRNA was added. The cells were cultured for another 36 hours before harvesting. As a negative control, dsRNA targeting the *AcGFP* gene was used. The templates for dsRNA synthesis were generated by PCR amplification from S2 cell genomic DNA using the following primers:

*uev1a*#1-F: GAATTAATACGACTCACTATAGGGAGAACGGAATTTCCGCTTACTG

*uev1a*#1-R: GAATTAATACGACTCACTATAGGGAGACGGACCGATGATCATGCC

*uev1a*#2-F: GAATTAATACGACTCACTATAGGGAGACACTAAAGATCGAGTGCG

*uev1a*#2-R: GAATTAATACGACTCACTATAGGGAGATGCCAGCTTCAGGTTCTC

*ben*#1-F: GAATTAATACGACTCACTATAGGGAGACCACGTCGCATCATCAAG

*ben*#1-R: GAATTAATACGACTCACTATAGGGAGAAGTCTTCGACGGCATATTTC

*ben*#2-F: GAATTAATACGACTCACTATAGGGAGACAGATCCGGACCATATTG

*ben*#2-R: GAATTAATACGACTCACTATAGGGAGATCAGTCTTCGACGGCATATTTC

*cdc27*#1-F: GAATTAATACGACTCACTATAGGGAGATCGCCCAGGATCTGATTAAC

*cdc27*#1-R: GAATTAATACGACTCACTATAGGGAGAGCAGCGACAGATCCTTCTTC

*cdc27*#2-F: GAATTAATACGACTCACTATAGGGAGAGATGATGGGCAAAAAGCTAAAG

*cdc27*#2-R: GAATTAATACGACTCACTATAGGGAGACCATCGGCCGATTGTTTC

*AcGFP*-F: GAATTAATACGACTCACTATAGGGAGATGCACCACCGGCAAGCTGCCTG

*AcGFP*-R: GAATTAATACGACTCACTATAGGGAGAGGCCAGCTGCACGCTGCCATC

### Human cell culture

The human colon cancer cell lines SW480 and HCT116, as well as the human embryonic kidney cell line 293T, were purchased from the American Type Culture Collection (ATCC) and cultured in Dulbecco’s Modified Eagle Medium supplemented with 10% FBS. The cultures were maintained at 37°C with 5% CO_2_ in a humidified incubator.

### Gene knockdown assay

Short hairpin RNA (shRNA) constructs were generated using the lentiviral vector *pLKO.1-CMV-copGFP-PURO*. Lentiviral particles were produced by co-transfecting each shRNA plasmid together with the packaging plasmids *psPAX2* and *pMD2.G* into 293T cells using Lipofectamine 2000, following the manufacturer’s protocol. SW480 and HCT116 cells were subsequently transduced with the harvested lentiviral supernatants to establish stable cell lines expressing *shNC* (negative control), *shUBE2V1*, or *shUBE2V2*. The following shRNA sequences were used:

*shNC*: CCTAAGGTTAAGTCGCCCTCG

*shUBE2V1 #1*: CTCGGGCAGATGACATGAAAT

*shUBE2V1 #2*: GCATCACCACAGGCTGGCTCA

*shUBE2V2 #1*: GTCTTAAATCAACAACCTTCT

*shUBE2V2 #2*: GCTCCTCCGTCAGTTAGATTT

### Overexpression assay

The lentiviral vector *pCDH-CMV* was used for gene overexpression, while *pMD2.G* and *psPAX2* served as lentiviral packaging plasmids. Target plasmids, *pCDH-CMV-UBE2V1* and *pCDH-CMV-UBE2V2*, were synthesized by the GENEWIZ company (Suzhou, China). Plasmids were transfected into 293T cells using Lipofectamine 2000 to generate lentiviral particles, according to the manufacturer’s protocol. SW480 and HCT116 cells were then transduced with these lentiviruses to establish stable cell lines expressing UBE2V1 and UBE2V2, respectively. To select for stably transduced cells, the cultures were maintained in medium containing 2 µg/mL puromycin.

### Quantitative real-time PCR (qRT-PCR)

Total RNA was extracted using the TriQuick Reagent kit and reverse-transcribed into cDNA using the SPARKscript II All-in-one RT SuperMix kit. qRTIZPCR was performed using the SYBR Green Premix Pro Taq HS qPCR kit. *GAPDH* was used as the endogenous control for normalization. The relative mRNA levels of target genes were determined using the 2^−ΔΔ*C*T^ method. The primer sequences used were:

*GAPDH*-F: ACAACTTTGGTATCGTGGAAGG

*GAPDH*-R: GCCATCACGCCACAGTTTC

*UBE2V1*-F: CGGGCTCGGGAGTAAAAGTC

*UBE2V1*-R: AGGCCCAATTATCATCCCTGT

*UBE2V2*-F: TGGACAGGCATGATTATTGGGC

*UBE2V2*-R: CTAACACTGGTATGCTCCGGG

### EdU incorporation assay

Cells were seeded in 24-well plates at a density of 100,000 cells per well. Cell proliferation was assessed using the EdU incorporation assay kit, following the manufacturer’s instructions. Briefly, EdU reagent was added to the culture medium, and cells were incubated for 2 hours to label DNA-synthesizing cells. After incubation, the medium containing EdU was removed, and cells were washed with PBS, fixed with fixative solution for 10 minutes, and then subjected to a click reaction for fluorescence labeling. Fluorescence microscopy was used to capture images and calculate the percentage of EdU^+^ cells.

### Colony formation assay

Cells were seeded in 6-well plates at a density of 500 cells per well and cultured at 37°C in a 5% CO_2_ incubator for 10 days without medium change. After 10 days, cells were gently washed with PBS to remove non-adherent cells, then stained with 0.5% crystal violet solution (containing 10% methanol and 1% acetic acid) for 15 minutes. The plates were subsequently rinsed with water to remove excess stain and air-dried. Colonies were photographed and counted using image analysis software.

### CCK8 cell viability assay

Cells were seeded in 96-well plates at a density of 2,000 cells per well. Cell viability was assessed over five days using a CCK8 kit following the manufacturer’s instructions. Measurements were taken on Day 0 (baseline) and daily from Day 1 to Day 4. At each time point, 10 µL of CCK8 solution was added to each well, and the plates were incubated for 2 hours at 37°C in a humidified incubator. Absorbance at 450 nm was measured using a microplate reader to assess cell viability and proliferation dynamics throughout the experimental period.

### Western blotting

Cells were lysed with RIPA buffer containing protease inhibitors, and protein concentrations were determined using a BCA Protein Assay Kit. Protein samples (20 µg per lane) were separated by SDS-PAGE and transferred to PVDF membranes. Membranes were blocked with 5% non-fat milk and incubated overnight at 4°C with primary antibodies: mouse anti-β-actin (Abmart, #M20011) at 1:5,000, rabbit anti-α-Tubulin at 1:5,000, rabbit anti-UBE2V1 at 1:2,000, and mouse anti-UBE2V2 at 1:2,000. Following washes, membranes were incubated with HRP-conjugated secondary antibodies and developed using ECL. Band intensities were quantified by densitometry.

### Immunohistochemistry assay

Tissue sections were deparaffinized with xylene, rehydrated through a graded ethanol series, and subjected to antigen retrieval in citrate buffer (pH 6.0) at 98°C for 10 minutes.

Endogenous peroxidase activity was blocked using 3% hydrogen peroxide, and nonspecific binding was blocked with 5% BSA. Sections were incubated overnight at 4°C with primary antibodies: rabbit anti-CycA at 1:200 and rabbit anti-Ki67 at 1:2,000. After incubation with HRP-conjugated secondary antibodies, sections were developed using DAB, counterstained with hematoxylin, and mounted. Images were captured using a light microscope.

### Animal experiment

Male *Balb/c* nude mice (7-week-old) were used to assess the tumorigenic potential of SW480 colon cancer cells. The mice were housed under specific pathogen-free conditions with a 12-hour light/dark cycle and had ad libitum access to food and water. SW480 cells were stably transfected with either the empty vector *pCDH-CMV* (control), *pCDH-CMV-UBE2V1* (UBE2V1-OE), or *pCDH-CMV-UBE2V2* (UBE2V2-OE), and 5×10^6^ cells were subcutaneously injected into the right flank of each mouse. Each group consisted of six mice. Tumor growth was monitored approximately one week after injection. Tumor volumes were measured every four days using calipers and calculated using the formula: volume = length × width^2^/2. The experiment was terminated when tumors reached ∼1,000 mm³, and mice were humanely euthanized according to institutional and national ethical guidelines.

### Image collection and processing

Fluorescent images were captured using a Zeiss LSM 710 confocal microscope (Carl Zeiss AG, BaWü, GER) and processed with Adobe Photoshop 2022 (San Jose, CA, USA), ImageJ (NIH, Bethesda, MD, USA), and ZEN 3.0 SR imaging software (Carl Zeiss).

## Acknowledgements

We gratefully thank Eric H. Baehrecke, Michael Buszczak, Zheng Guo, Yuu Kimata, Ruth Lehmann, Erika Matunis, Duojia Pan, Zhaohui Wang, Wei Xiao, Addgene, ATCC, BDSC, CEMCS, DGRC, GenBank, and THFC for providing antibodies, plasmids, cell lines, and fly strains. This study was supported by National Natural Science Foundation of China (NSFC) grants to Shaowei Zhao (32270841, 32070871), Shian Wu (32170714), and Hongru Zhang (32400759), as well as by a Natural Science Foundation of Tianjin grant (S24ZDD020) to Hongru Zhang.

## Author contributions

S.Z. conceived the project. S.Z., S.W., and H.Z. supervised the studies. Q.Z., Yun.W., X.F., Z.W., Y.Z., L.Y., Yue.W., M.Y., D.S., and R.Z. performed the experiments. S.Z. wrote the manuscript and all authors commented on it.

---

**Figure 2-figure supplement 1.**
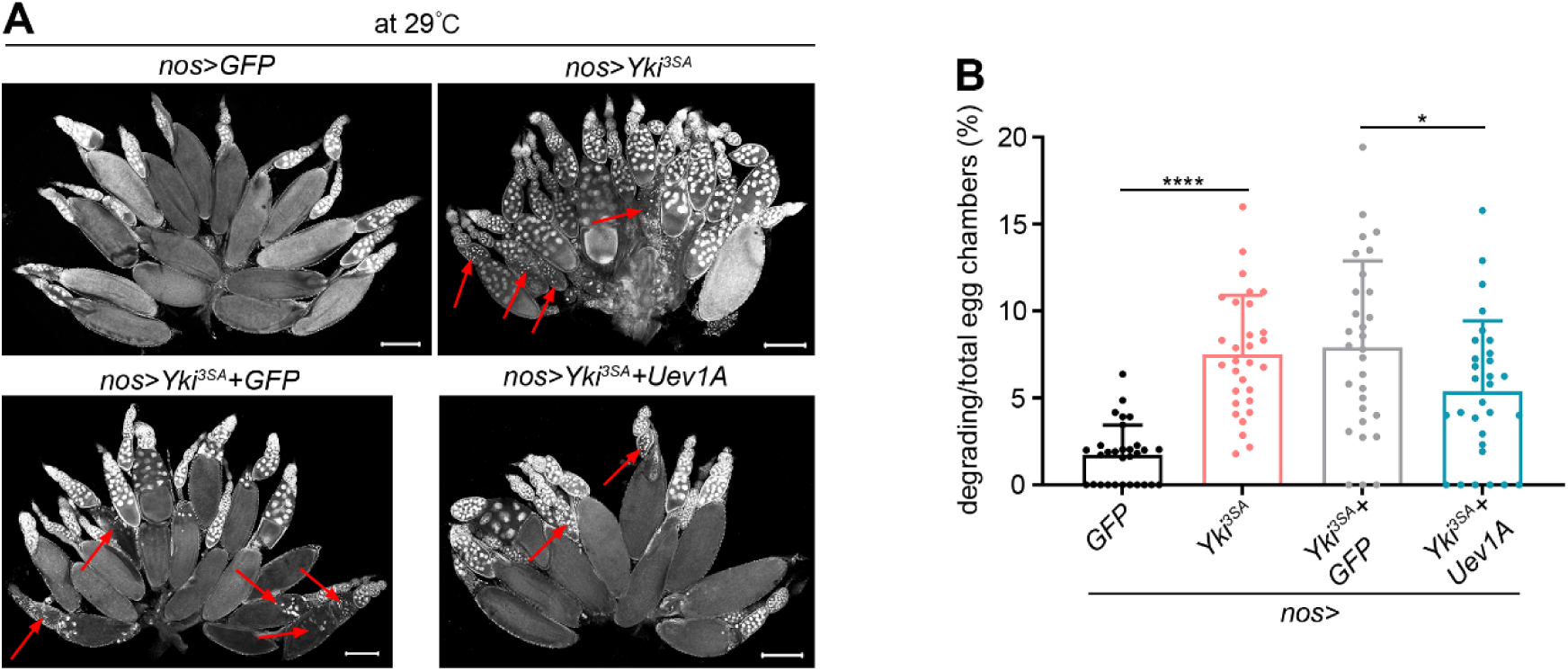
Uev1A protects against *Yki^3SA^*-induced nurse cell death. (A) Representative ovaries (DAPI staining). The red arrows denote degrading egg chambers. Scale bars: 200 μm. (B) Quantification data. 30 ovaries from 7-day-old flies were quantified for each genotype. Statistical significance was determined using t test: * (*P* < 0.05) and **** (*P* < 0.0001). See also **Source data 2 and 3**.

**Figure 3-figure supplement 1.**
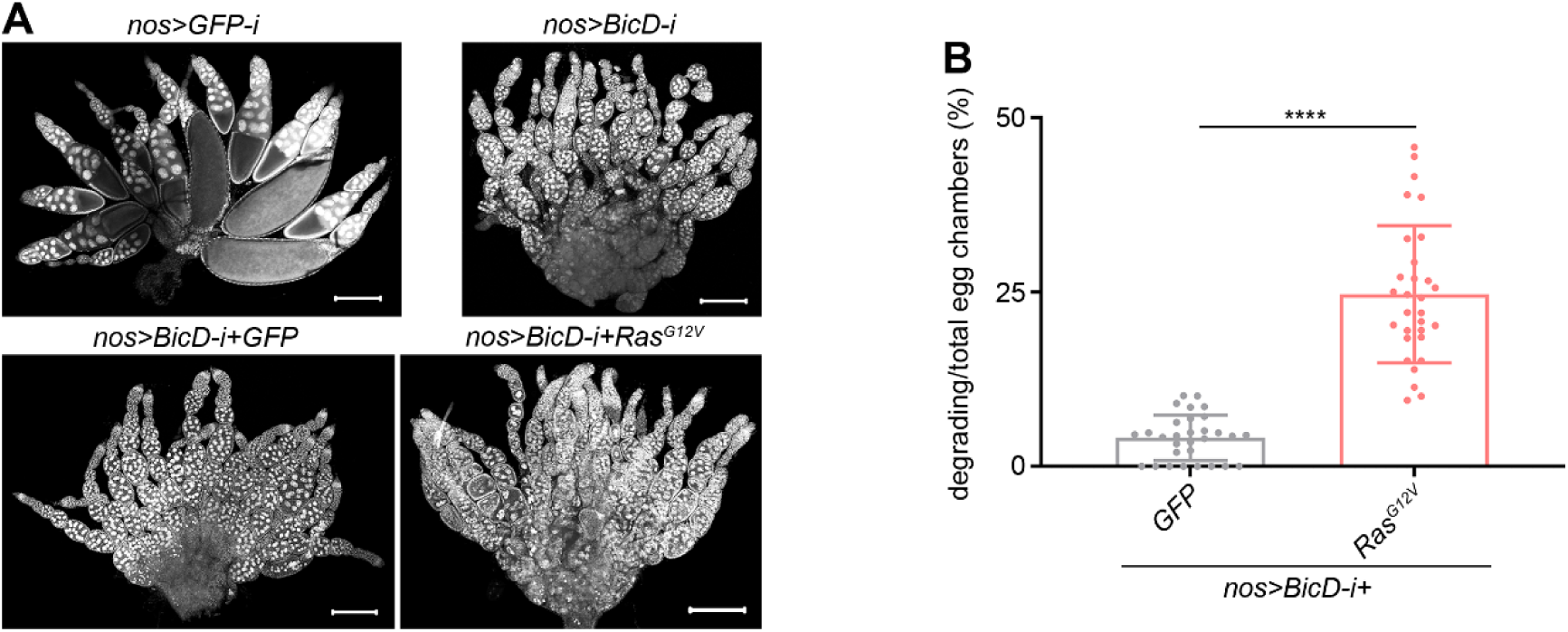
Oncogenic *Ras^G12V^* intrinsically triggers nurse cell death. (A) Representative ovaries (DAPI staining). Scale bars: 200 μm. (B) Quantification data. 30 ovaries from 3-day-old flies were quantified for each genotype. Statistical significance was determined using t test: **** (*P* < 0.0001). See also **Source data 2 and 3**.

**Figure 3-figure supplement 2.**
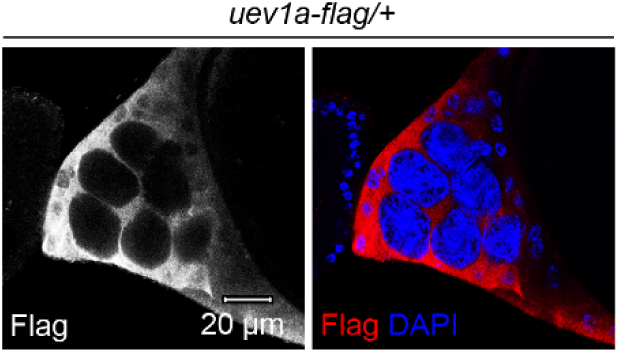
Uev1A is expressed in stretch follicle cells. Representative ovaries.

**Figure 3-figure supplement 3.**
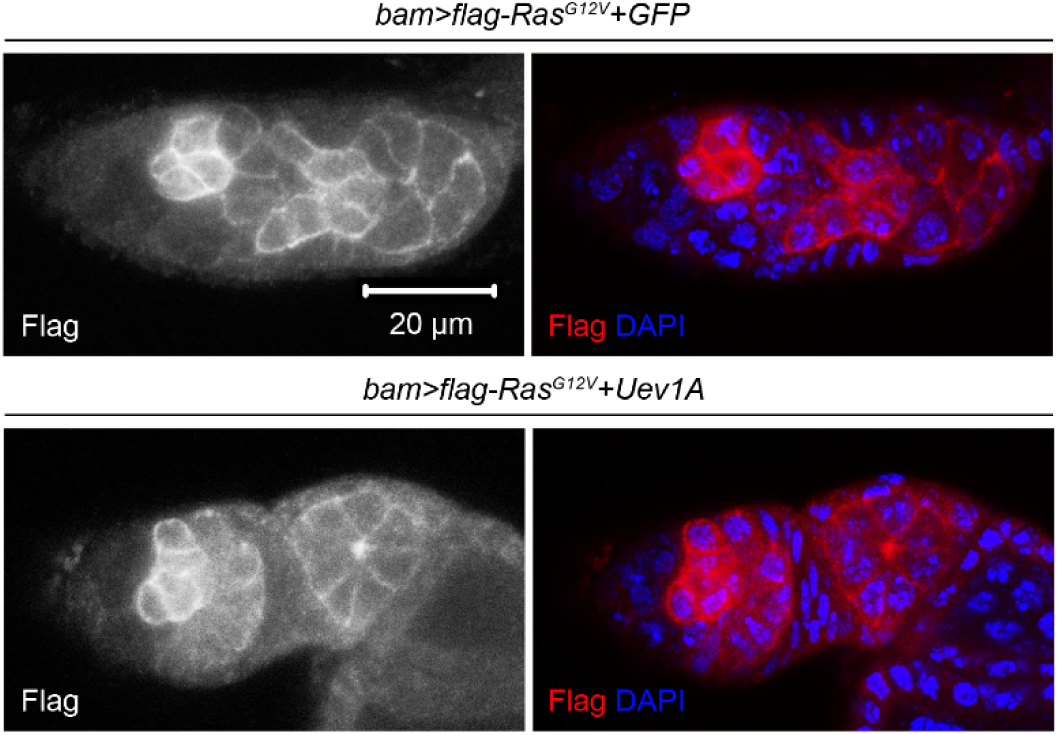
Uev1A does not directly degrade the Ras^G12V^ oncoproteins. These experiments were performed at 29°C. All images are of the same magnification. See also **Source data 2.**

**Figure 5-figure supplement 1.**
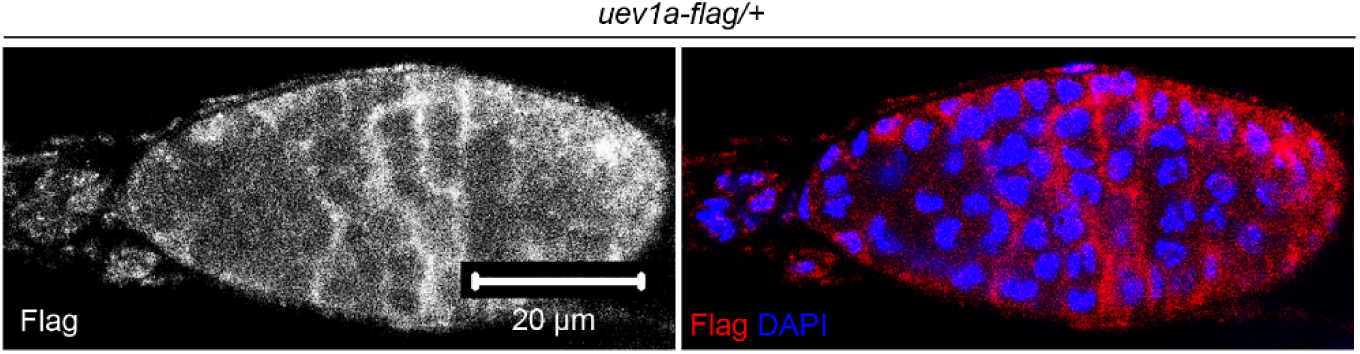
Expression pattern of Uev1A in germarium. Representative sample. See also **Source data 2.**

**Figure 5-figure supplement 2.**
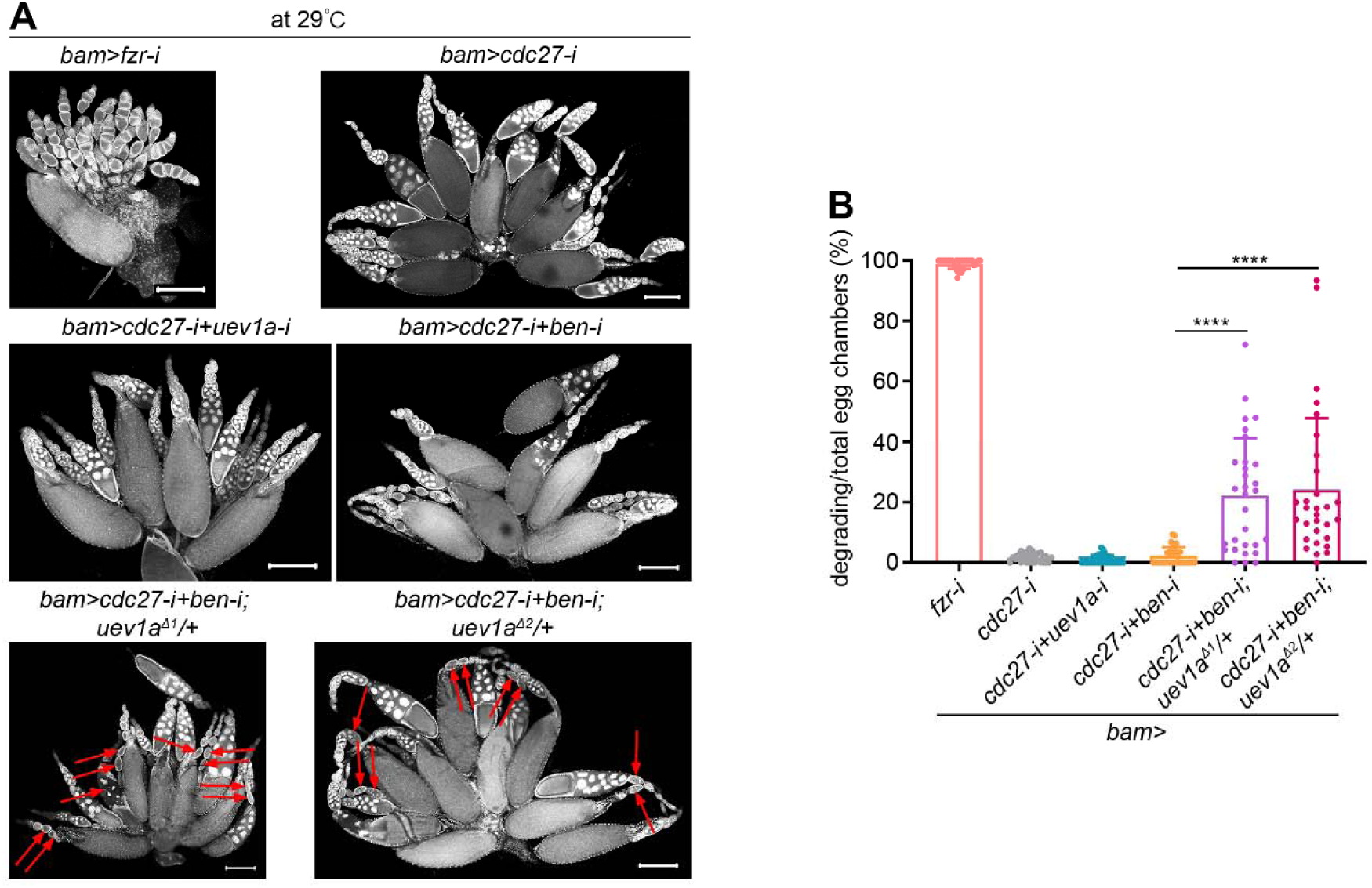
Uev1A, Ben, and Cdc27 work together to protect nurse cells from death during normal oogenesis. These experiments were performed at 29°C. (A) Representative ovaries (DAPI staining). The red arrows denote degrading egg chambers. Scale bars: 200 μm. (B) Quantification data. 30 ovaries from 7-day-old flies were quantified for each genotype. Statistical significance was determined using one-way ANOVA: **** (*P* < 0.0001). See also **Source data 2 and 3**.

**Figure 6-figure supplement 1.**
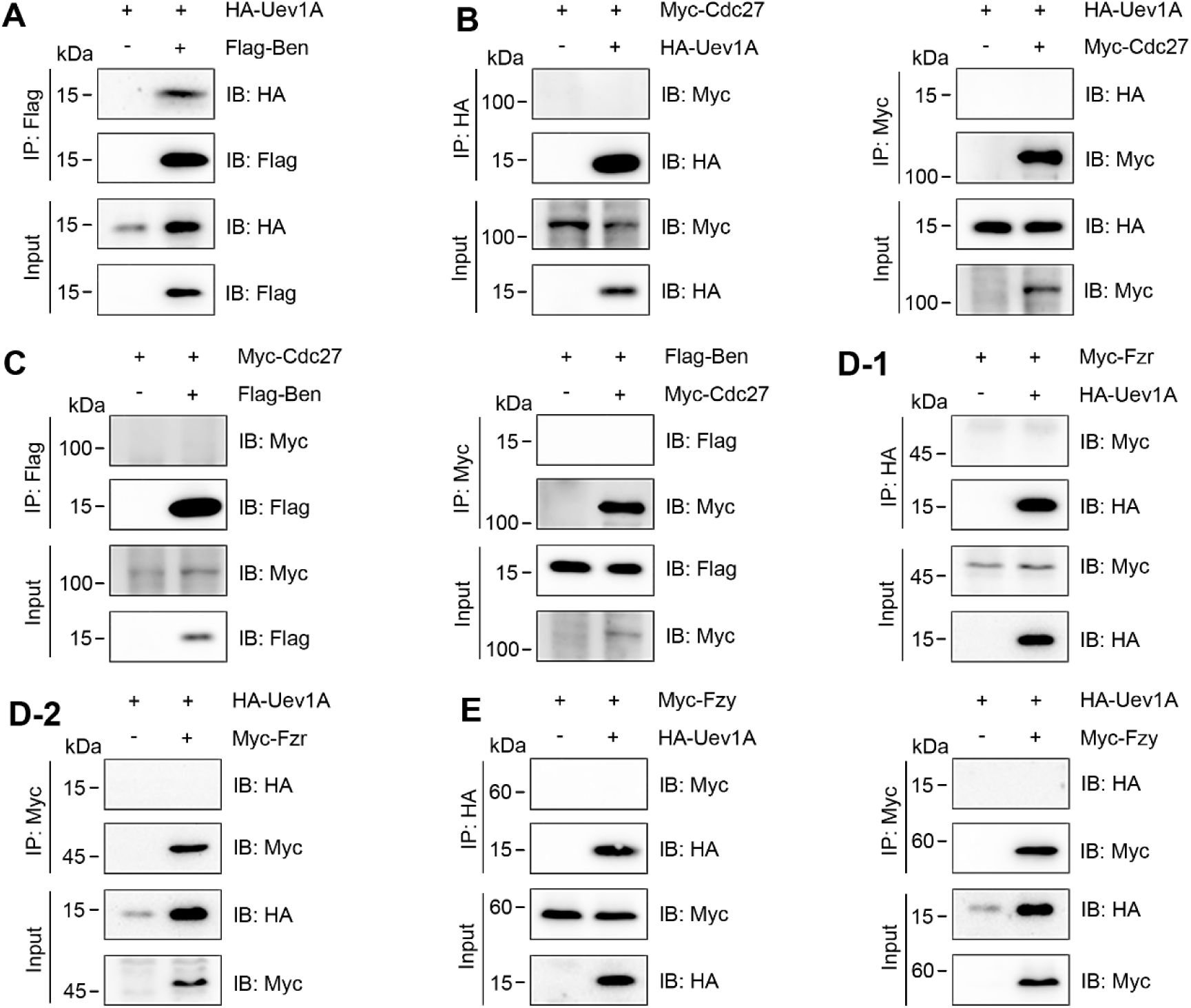
Co-IP results. The tagged proteins were co-expressed in S2 cells to assess physical interactions. Physical interaction was observed between Uev1A and Ben (A). No interaction was detected between Uev1A and Cdc27, Fzr, or Fzy (B, D, E), nor between Ben and Cdc27 (C).

**Figure 6-figure supplement 2.**
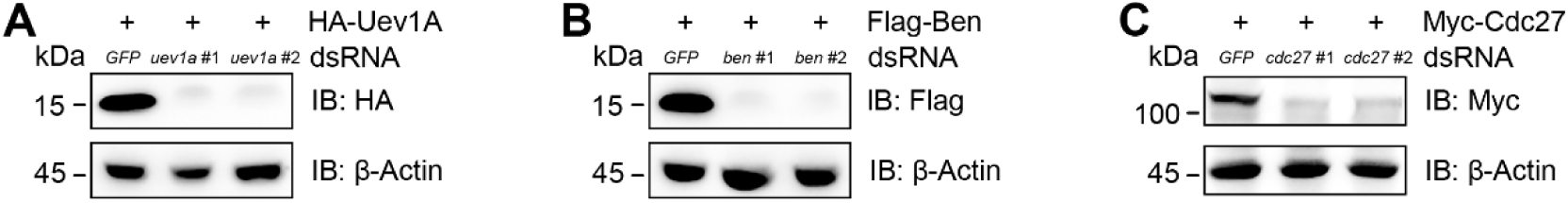
RNAi efficiency assays. The #1 dsRNAs were employed in RNAi assays targeting *uev1a*, *ben*, and *cdc27* in Figure 6C, D, and F.

**Figure 8-figure supplement 1.**
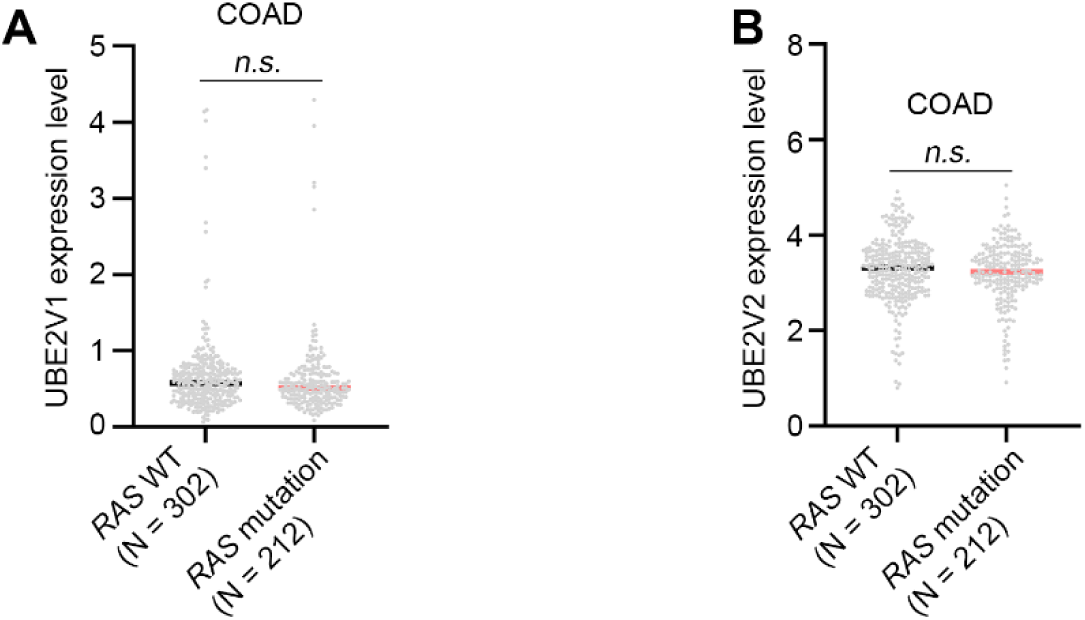
TCGA analysis comparing UBE2V1 and UBE2V2 expression levels in colorectal cancer patients with and without *RAS* mutations. The TCGA patient data were downloaded from the UCSC Xena website: the RNA-seq data (Version: 05-09-2024) and the somatic mutation data (Version: 08-05-2024). Statistical significance was determined using t test: *n.s.* (*P* > 0.05).

**Figure 8-figure supplement 2.**
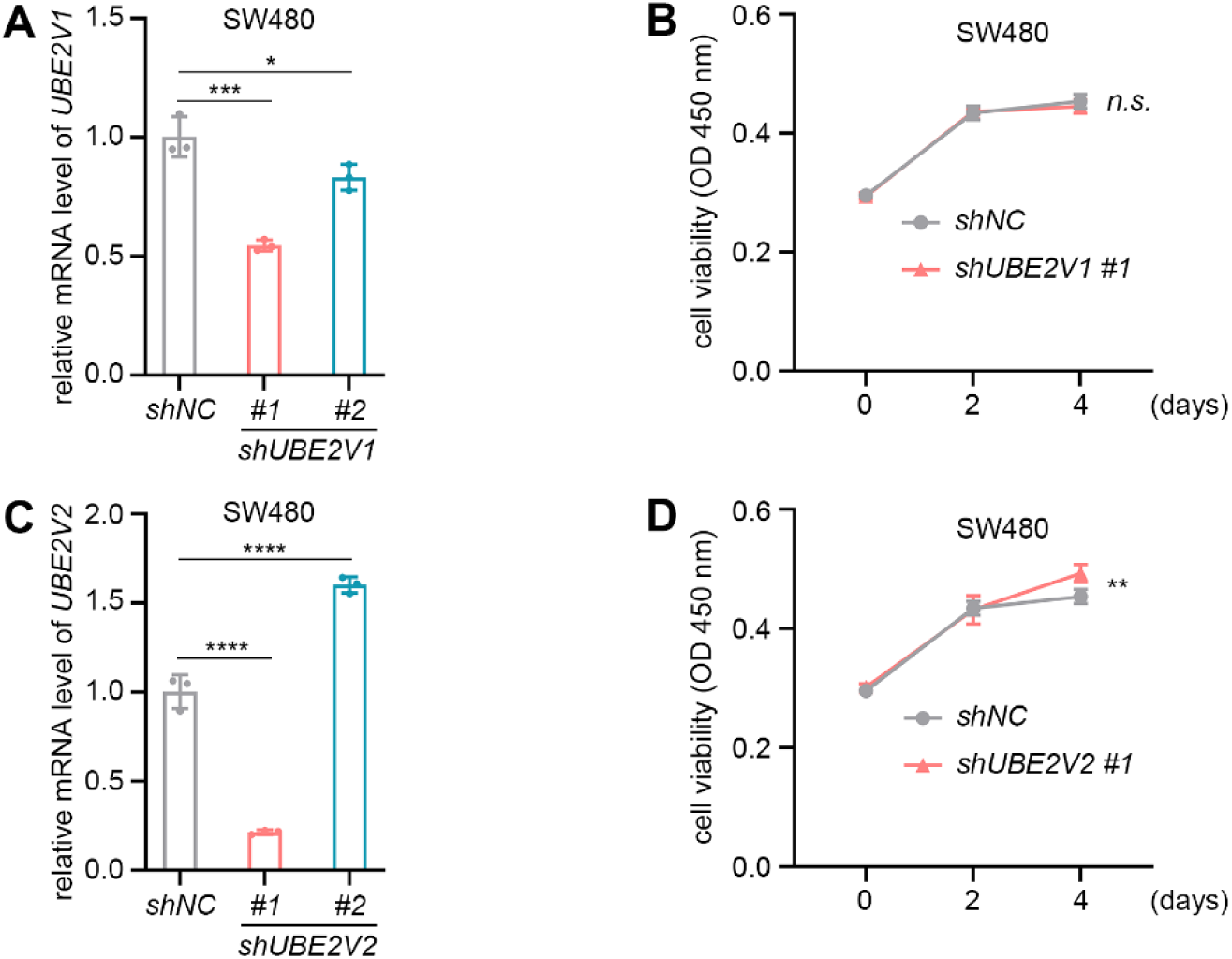
Knocking down either *UBE2V1* or *UBE2V2* alone mildly influences the growth of colorectal cancer cell lines. (A and C) The knock-down efficiency assays. The relative mRNA levels were normalized to *GAPDH*. Three independent replicates were conducted, and statistical significance was determined using one-way ANOVA. (B and D) Assays to evaluate the effects of *UBE2V1-* and *UBE2V2-RNAi* on colony formation and cell viability in SW480 cells. Six independent replicates were conducted at each time point, and statistical significance was determined using two-way ANOVA with multiple comparisons. *n.s.* (*P* > 0.05), * (*P* < 0.05), ** (*P* < 0.01), *** (*P* < 0.001), and **** (*P* < 0.0001).

**Figure 9-figure supplement 1.**
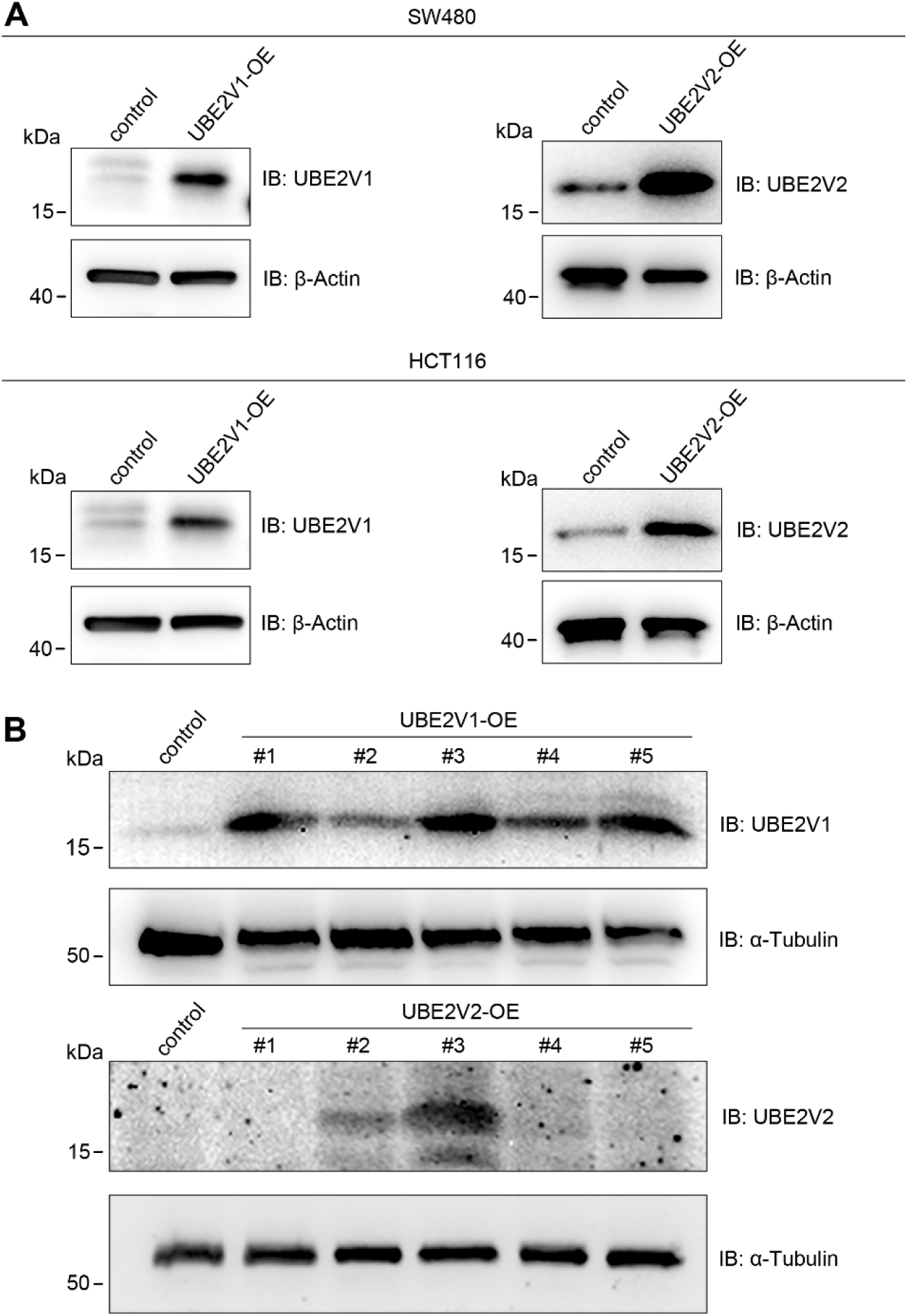
Validation of UBE2V1/2 overexpression in colorectal cancer cell lines. (A) Western blotting to confirm the transient overexpression of UBE2V1 and UBE2V2 in SW480 and HCT116 cell lines. β-Actin was used as the loading control. (B) Western blotting to confirm the stable overexpression of UBE2V1 and UBE2V2 in SW480 cells, with UBE2V1-OE #1 and UBE2V2 #3 cell lines utilized in subcutaneous tumorigenesis assays. α-Tubulin was used as the loading control. In both (A) and (B), cells transfected with an empty overexpression vector served as the control.

**Figure 9-figure supplement 2.**
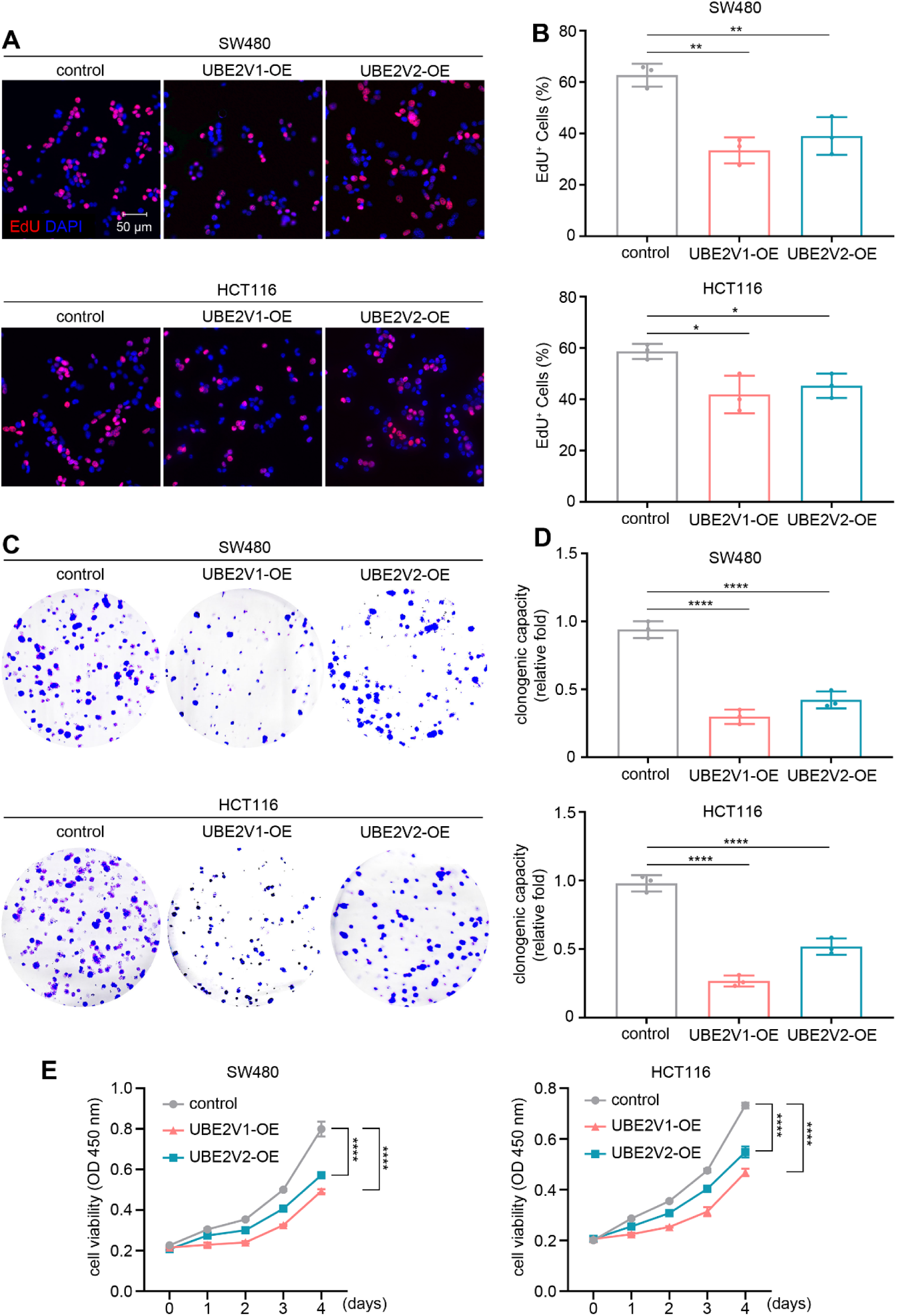
UBE2V1/2 overexpression suppresses the growth of colorectal cancer cell lines. (A and B) EdU incorporation assays to assess the effects of UBE2V1 and UBE2V2 overexpression (OE) on cell proliferation in SW480 and HCT116 cells. Empty OE vector was used as the control. All images in (A) are of the same magnification. (C-E) Assays to evaluate the effects of UBE2V1- and UBE2V2-OE on colony formation and cell viability in SW480 and HCT116 cells. In (B and D), three independent replicates were conducted, and statistical significance was determined using one-way ANOVA. In (E), five independent replicates were conducted, and statistical significance was determined using two-way ANOVA with multiple comparisons. * (*P* < 0.05), ** (*P* < 0.01), and **** (*P* < 0.0001).

